# An injectable soft implant for long-acting, reversible, ultra-stable release of therapeutics

**DOI:** 10.64898/2026.02.25.707913

**Authors:** Ceren Kütahya, Luca Panariello, Adrian Najer, Tatiana Rizou, André Shamsabadi, Giulia Brachi, David J. Peeler, Lukeriya Zharova, Thomas F. F. Fernandez Debets, Patrick Peschke, Anna P. Constantinou, Ruoxiao Xie, Yuxi Cheng, Ross Burdis, Alejandro Suarez-Bonnet, Martina Cihova, Jonathan Yeow, Fredrik Schaufelberger, Ilaria Malanchi, Molly M. Stevens

**Author notes:** **Corresponding Author** Correspondence to Molly M. Stevens.

## Abstract

Providing long-term (>6 months) zero-order drug release from easily administered formulations is a key challenge in improving patient adherence and facilitating access. Herein, we report the design and development of an injectable, biodegradable, long-acting polymeric microparticle-embedded hydrogel platform for prolonged, zero-order release of therapeutics. This “soft implant” is injectable for ease of administration and can be retrieved *via* a small incision, allowing for discontinuation of therapy if desired. Central to the platform are surface-eroding poly(orthoester) (POE) microparticles, which were molecularly tailored to tune zero-order drug release across a wide range of timeframes. We demonstrate the clinical potential of the “soft implant” using levonorgestrel, a contraceptive agent requiring sustained dosing. *In vitro*, we observed zero-order release for 300 days, projected for >12 months, with behavior consistent with surface erosion further supported through Raman chemical mapping. *In vivo* studies confirmed zero-order release for six months, projected to 12 months, from a subcutaneous injection in rats. We envision that our platform could transform therapies that require long-term, regular drug dosing, significantly improving compliance and therapy outcomes.

## Main

Long-term therapy, which typically demands frequent dosing, compromises patient adherence and limits therapeutic effectiveness^1, 2^. To address this challenge, long-acting formulations are designed to release drugs gradually from a single administration, maintaining therapeutic levels over extended periods^3, 4^. By providing sustained drug exposure, these platforms can enhance regimen efficacy, decrease side effects, improve adherence, and reduce healthcare burdens on society. The development of effective long-acting formulations is an unresolved challenge for many clinical scenarios, from treatment of chronic disorders (e.g. diabetes, cardiovascular diseases, cancer, etc.), relapsing-release disorders (e.g. inflammatory bowel disease, asthma), acute illnesses (e.g. tuberculosis, hepatitis B, etc.), and for prophylactic applications (e.g. contraception, HIV pre-exposure prophylaxis, vaccines, etc.)^4-6^. Technologies that can facilitate patient adherence to therapeutic regimes hold major promise in improving the health of the general population in an impactful and cost-effective way^2, 6^. While long-acting drug release devices covering prolonged release windows (>6 months) have been reported, they generally require invasive administration procedures (e.g. trocar) and oftentimes surgical removal at the end of the treatment^7^. Recently, significant progress has been achieved in the development of injectable long-acting release systems, including *in situ*-forming depots, often relying on the development of hydrophobic prodrugs with favorable pharmacokinetic (PK) profiles^8-10^ or leveraging the use of micronized crystals with slow dissolution rates^11^. These systems are difficult to generalize both in terms of the nature of the active component released and the length of the release window. Furthermore, they can also require the injection of large volumes of organic solvents, with associated side effects^4^.

The development of biodegradable polymeric formulations that mediate drug release is an attractive alternative for long-acting formulations, where polymer degradation is used as the driving parameter to control drug release, providing a platform system for the controlled release of multiple therapeutics. Through precise microfabrication, micron scale drug-loaded microparticles can be manufactured, providing an injectable platform for drug release^4, 12-15^. Current polymer-based injectable formulations for sustained drug release are mostly based on hydrolytically degradable polyesters, such as poly(lactic-*co*-glycolic acid) (PLGA) or polycaprolactone (PCL)^16, 17^. However, these formulations generally exhibit ‘irregular’ drug release profiles, often characterized by ‘bursts’ and ‘plateaus’ owing to their bulk-eroding profile, making them unsuitable for prolonged, dose-compliant drug release^18-20^. In contrast to PLGA, surface-eroding polymers (specifically poly(orthoesters), (POEs)) have been utilized to fabricate solid devices that promote the stable, prolonged linear (‘zero-order’) release of drugs as the matrix degrades.^21-26^ Achieving zero-order release and reduced initial burst is crucial to mitigate concentration-dependent side effects, associated for instance with glucagon-like peptide 1 receptor agonists for the treatment of type 2 diabetes, or with antiepileptic drugs. Effective long-acting delivery platforms using POEs have so far been limited to large implantable devices^25, 26^, which require invasive procedures for placement.

A further challenge of injectable long-acting release formulations is retrieving the platform if the patient requires treatment termination, particularly in cases involving unforeseen side-effects or development of drug resistance. This is also relevant for long-acting contraceptive systems, as the patient may desire to terminate use early to restore fertility. Examples of polymeric reversible long-acting formulations consist primarily of externally applied PLGA-based microneedle patches, which have demonstrated sustained release of levonorgestrel (LNG) for only 2 months in rats^27-29^.

Herein, we developed an injectable, bioerodible delivery platform offering long-term, sustained drug release with optional retrievability (**Fig. 1**). Using contraception as a case study, we demonstrated this platform with LNG as active compound. We designed and synthesized poly(orthoester) (POE) polymer matrices to fabricate monodisperse LNG-loaded microparticles *via* microfluidics and embedded these microparticles in a hydrogel carrier to enable non-invasive injection and retrieval. We demonstrate how sustained long-term, zero-order drug release can be tuned from 1 up to 12 months by controlling the chemical composition of POE polymers. This high level of control over zero-order release kinetics for up to a year is a significant advance for the field and is promising for the development of next-generation long-acting drug delivery systems.

**Fig. 1:**
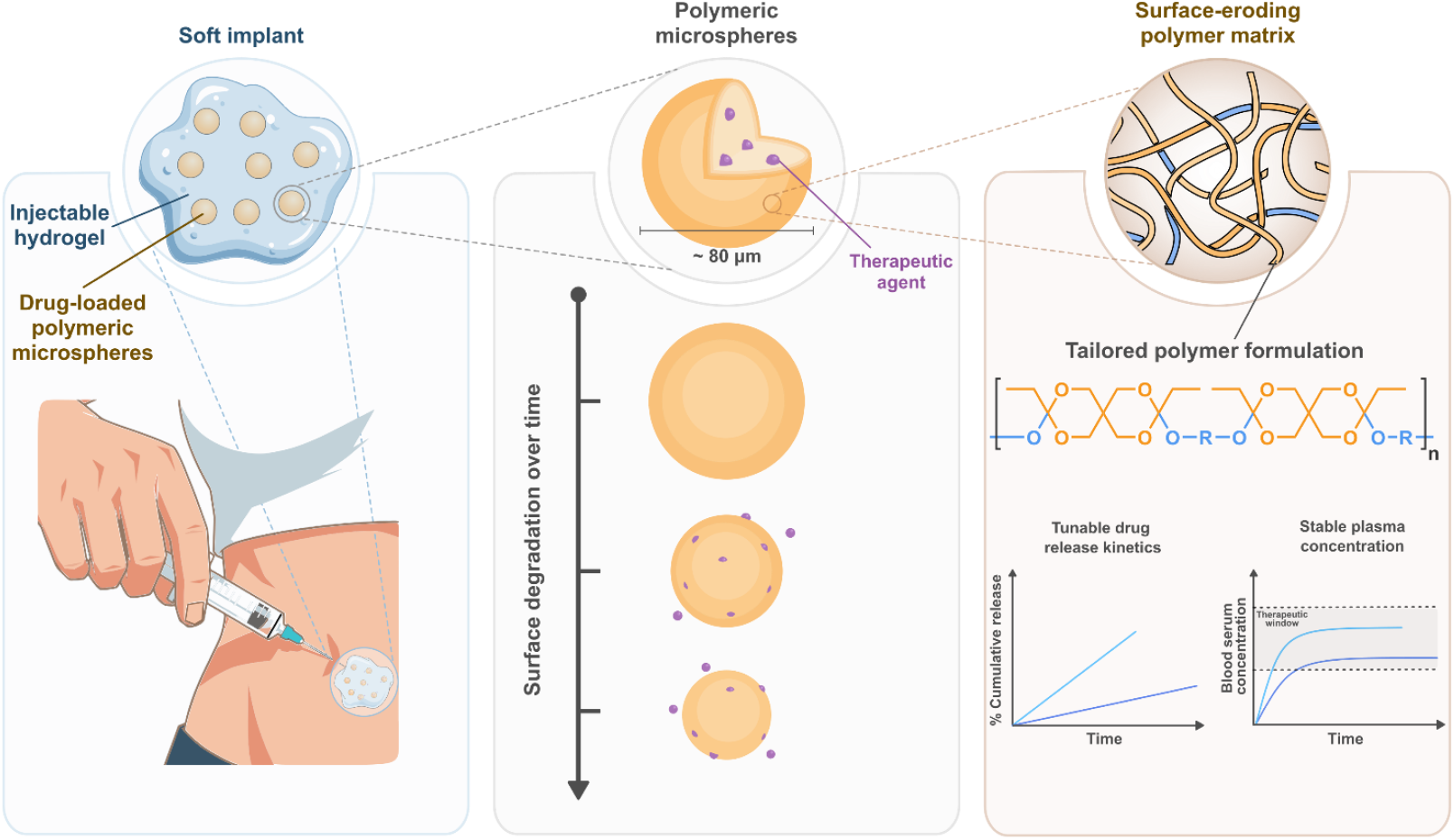
A modular, biodegradable, and injectable hydrogel platform embedded with polymeric microparticles for the tailored, zero-order release of therapeutics. The hydrogel composite can be easily administered through a 23G needle. Because the therapeutic agent is uniformly dispersed within homogeneous, surface-eroding microparticles, its release remains highly stable and exhibits zero-order kinetics. Furthermore, the specific release rate can be precisely tuned by tailoring the polymer composition whilst maintaining this zero-order profile.

## Results

### Poly(orthoester) library for steady-state release of LNG

Long-acting drug release systems are particularly relevant in the context of contraception, with a significant need for safe, effective, low-cost methods capable of sustained, long-term release. Long-acting contraceptives are indispensable in addressing issues of patient compliance and accessibility, particularly in low-income countries and underserved communities^30^, providing women with greater reproductive autonomy. Although existing long-acting reversible contraceptive (LARC) delivery platforms are highly reliable for preventing pregnancy, these methods are invasive, requiring either the physical insertion of an intrauterine device (IUD) or an implant *via* subdermal incision. Currently, progesterone-based injectable contraceptives exist as a non-invasive platform only capable of providing protection for only 12-14 weeks^31^, whereas oral contraceptives offer short-term efficacy and are notably susceptible to patient non-compliance^32^.

To realize a drug delivery platform that exhibits zero-order release of the contraceptive LNG, we sought a highly modular surface-erodible polymer platform. POEs are a class of rationally designed polymers known to display surface erosion *via* hydrolysis, generating monomeric diols and a trace amount of acid^33^. POEs were synthesized through the nucleophilic addition of diols to diketene acetal (**Table 1**). Bulk erosion in these polymers is inhibited owing to their hydrophobic nature, which restricts water penetration through the bulk material^33, 34^. To increase the hydrolysis rate of POE materials, latent acid (lactic acid or glycolic acid) or acidic excipients were incorporated^26, 33, 35-37^ to promote auto-catalytic hydrolysis of the orthoester backbone. Conversely, basic excipients have been included in the polymer matrix to inhibit hydrolysis, stabilizing the bulk polymer and further facilitating erosion to primarily occur at the outer surface^21, 25, 38^.

**Table 1:** Polymers used to formulate microparticles for tunable LNG release kinetics^a^.

| Polymer | H:D:T:G:C:P<br>(mol%) | $M_w^d$<br>(kDa) | $M_n^d$<br>(kDa) | $\bar{D}^d$ |
| --- | --- | --- | --- | --- |
| POE-C9G1 | 0:0:0:10:90:0 | 8.5 | 6.7 | 1.27 |
| POE-H | 100:0:0:0:0:0 | 3.6 | 2.0 | 1.77 |
| POE-C | 0:0:0:0:100:0 | 4.2 | 3.0 | 1.41 |
| POE-H1C2 | 33:0:0:0:67:0 | 7.7 | 4.0 | 1.92 |
| POE-D1C2 | 0:33:0:0:67:0 | 6.0 | 4.0 | 1.50 |
| POE-C9T1 | 0:0:10:0:90:0 | 6.9 | 4.1 | 1.67 |
| POE-H3C6P1 | 30:0:0:0:60:10 | 4.8 | 3.5 | 1.35 |
| POE-H3C6T1 | 30:0:10:0:60:0 | 4.0 | 3.0 | 1.35 |
| Resomer® RG 756 S<sup>b</sup> | N/A | 76-115 | - | - |
| Resomer® RG 503 H<sup>c</sup> | N/A | 24-38 | - | - |

<sup>a</sup>Conditions: All polymerizations were conducted using DETOSU at a concentration of 50 mol% relative to the total amount of diols.
<sup>b</sup>Commerical Resomer® RG 756 S, (poly(D,L-lactide-co-glycolide) ester terminated, lactide:glycolide 3:1 molar equivalent).
<sup>c</sup>Commerical Resomer® RG 503 H, (poly(D,L-lactide-co-glycolide) acid terminated, lactide:glycolide 1:1 molar equivalent).
<sup>d</sup>Determined by gel permeation chromatography using dimethylformamide as the eluent and polymethyl methacrylate standards.

For the realization of a polymer matrix demonstrating a long-term degradation profile, we hypothesized that the reduced quantity of orthoester linkages in linear POE structures, compared to networked POE formulations synthesized utilizing polyols, would inhibit the erosion rate and extend concomitant drug release. Additionally, we proposed that the degradation kinetics and physical properties of POE structures could be systematically fine-tuned by diol monomer chemistry. By investigating the structure-function relationship of POE materials, we aimed to synthesize compositions that yielded defect-free MPs with the desired topology and degradation profiles. With this in mind, a library of POE formulations was prepared utilizing 3,9-diethylidene-2,4,8,10-tetraoxaspiro(5.5)undecane (DETOSU) as the diketene acetal and a variety of diols— 1,6-hexanediol (**H**), 1,10-decanediol (**D**), *trans-*1,4-cyclohexanedimethanol (**C**), triethylene glycol (**T**), glycerol monomethacrylate (**G**), and 1,4-bis(2-hydroxyethyl)piperazine (**P**)— at particular ratios (**Table 1**). As a control, commercially sourced PLGA, known for its bulk-eroding behavior^39, 40^, was used to benchmark the performance of the experimental polymers.

### Microfluidic fabrication of highly uniform polymeric MPs that readily integrate in a hydrogel carrier

To provide a scalable and modular platform, we adopted a microfluidic emulsification approach for MP production. The use of microfluidics is crucial to produce defect-free MPs with highly uniform sizes and shapes to achieve quantitative drug encapsulation and zero-order release kinetics. Furthermore, size control is critical to ensure injectability of the final platform, as even a small fraction of particles with size larger than the internal needle diameter could lead to clogging. Additionally, this manufacturing technique is compatible with upscaling production through parallel, continuous flow synthesis^41^. For our custom-made microfabrication setup, the microfluidic chips were fabricated using a device with coaxially aligned glass microcapillaries^42^. The configuration consists of a flow-focusing device where an aqueous and an organic phase (water-immiscible, containing polymer, excipients and LNG 5% (w/w) unless otherwise stated) flow through a capillary in a counter-flow pattern with droplets pinching at the capillary inlet (**Fig. 2a**), producing a stable emulsion of monodisperse droplets. The droplets were collected in petri dishes, and the organic solvent was allowed to evaporate overnight yielding solid polymeric microparticles. This microfabrication method can be employed to produce MPs of the polymers in **Table 1** with fixed particle diameter of ∼80 µm (observed by scanning electron microscopy (SEM), **Fig. 2b-f and Supplementary Fig. 1-2**), hence allowing the assessment of different formulations without size-related variability. SEM images of cross-sections demonstrate how these particles exhibit a structure with minimal internal porosity (**Fig. 2e**).

**Fig. 2:**
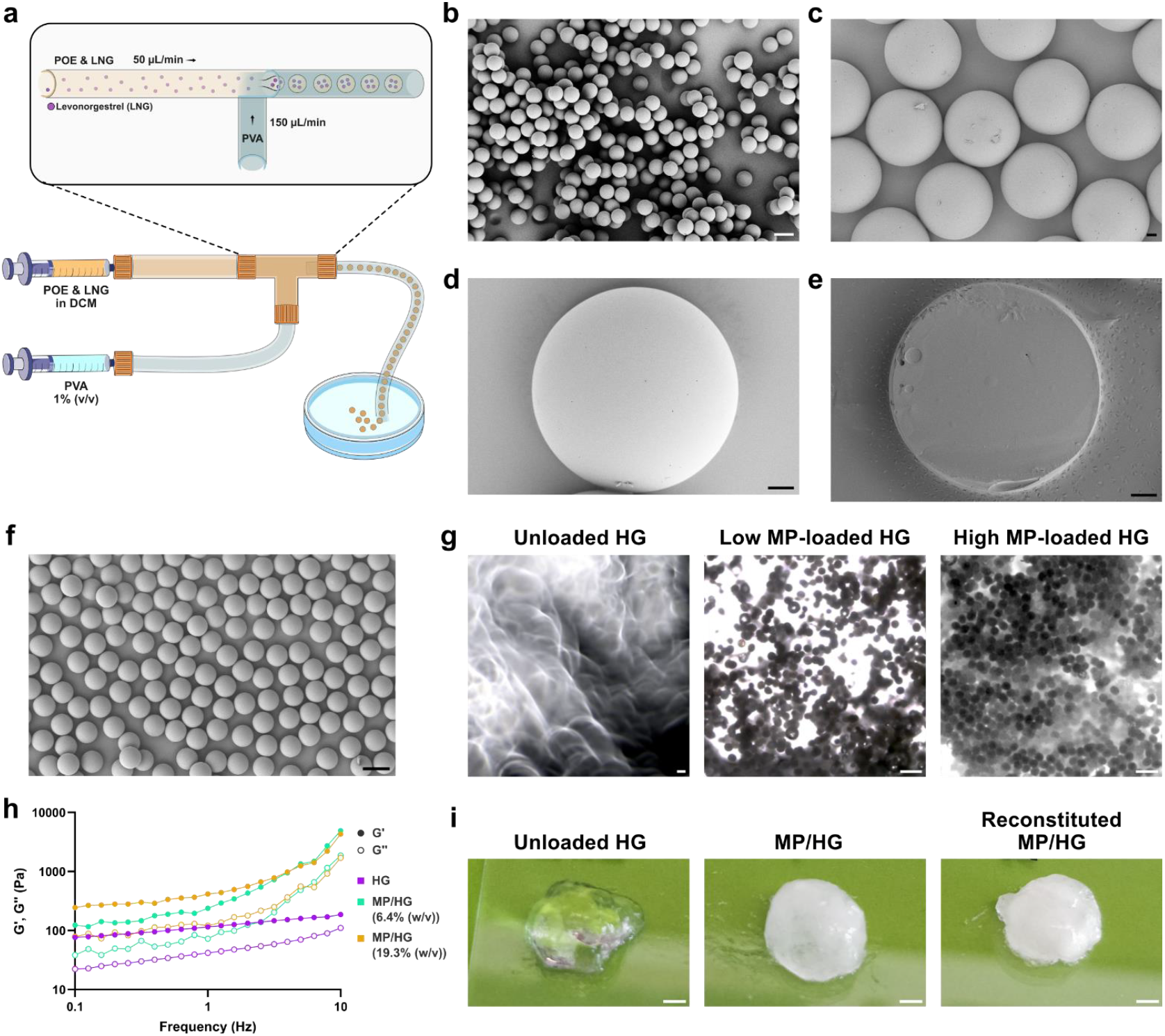
Fabrication of polymeric microparticle/hydrogel composite. **a**, Schematic of the custom-made microfabrication set-up with droplets formed when entering the smaller capillary. **b**,**c**, SEM images of POE-H3C6P1 (LNG 5% (w/w)) MPs. Scale bars: 100 µm (b), 10 µm (c). **d**,**e**, SEM images of the MP cross-section. Scale bar: 10 µm. **f**, SEM image of PLGA (Resomer^®^ RG 503 H) (LNG 5% (w/w)) MPs. Scale bar: 100 µm. **g**, Representative bright-field microscopy images of the unloaded HG and PLGA MPs loaded HG (6.4% (w/v) and 19.3% (w/v)). Scale bar: 200 μm. **h**, Frequency sweep of unloaded HG and PLGA MPs (6.4% (w/v) and 19.3% (w/v)) loaded HG. **i**, Pictures of HG, PLGA MPs embedded (19.3% (w/v)) and reconstituted formulation. Scale bar: 2 mm.

While microfluidic fabrication ensures reproducible and uniform MP size well below the inner diameter of a 23G needle, effectively injecting microparticles through conventional syringes remains challenging, often requiring the use of additives and custom-designed syringes to ensure injectability^43^. Furthermore, microparticles injected on their own are non-retrievable, precluding elective treatment discontinuation. Considering these limitations, we employed medical-grade hyaluronic acid (HA) dermal fillers (Belotero^®^ fillers with a HA content of 2.5% (w/v)) as an injectable and biodegradable polymer hydrogel (HG) as a carrier for the MPs. These dermal fillers are approved for use in clinical applications as injectable treatments for up to 18 months^44, 45^. Initially, we evaluated the injectability of our platform using HG embedded with ∼80 µm PLGA MPs (desired size for our platform, **Fig. 2f**) at low (6.4% (w/v)) and high (19.3% (w/v)) particle concentrations (**Fig. 2g**). Rheological characterization of MP/HG composites was carried out at 37 °C. A frequency sweep test confirmed the presence of an elastic HG network, with increased G’ and G’’ with increasing MP concentration in the matrix (**Fig. 2h**). To ensure the stability and homogeneity of the platform prior to administration, a facile reconstitution process was developed by freeze-drying MP/HG composites and re-hydrating before injection. Reconstituted MP/HG formulations showed similar appearance and particle distributions to the original fabrication method (**Fig. 2i**).

### POE microparticles-hydrogel composites exhibit zero-order sustained release with controllable release windows

*In vitro* release experiments from POE MPs were performed by incubating MP/HG composites in PBS containing 3% (w/v) 2-hydroxypropyl-β-cyclodextrin and 0.1% (w/v) NaN_3_ at 37 °C. **Fig. 3** and **Extended data Fig. 1** show the cumulative release of LNG from MPs formed with different POE formulations (Table 1) containing 5% (w/w) LNG (unless otherwise stated). We observed that most POE formulations provided zero-order release with release rates strongly dependent on the monomer composition of the polymer backbone.

**Fig. 3:**
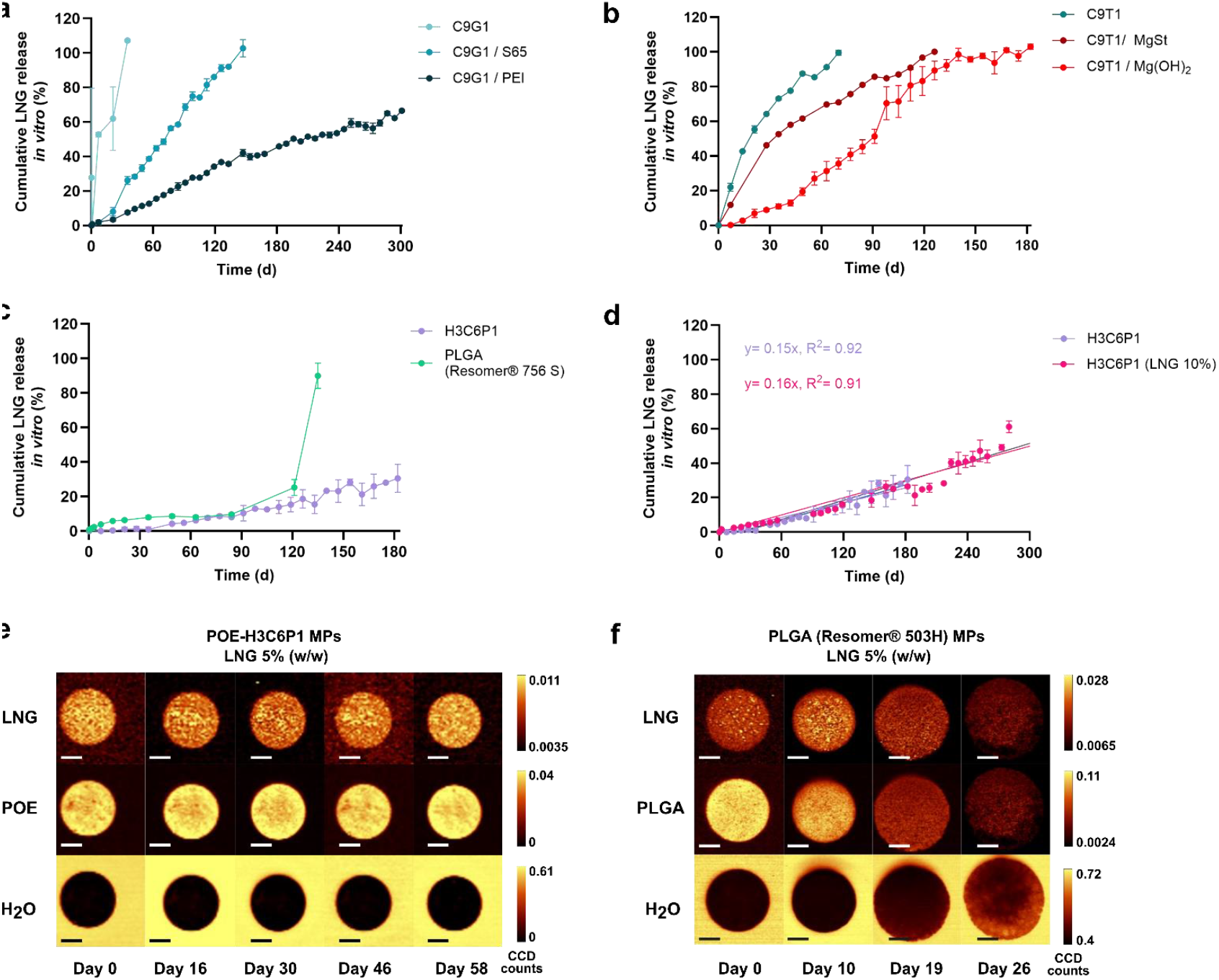
*In vitro* characterization of LNG-loaded POE microparticles (MPs). **a–d**, Cumulative *in vitro* release of LNG from POE MPs incubated in PBS containing 3% (w/v) 2-hydroxypropyl-β-cyclodextrin and 0.1% (w/v) sodium azide at 37 °C, plotted over time. Data represents mean ± s.d. (n = 3 different release reservoirs). **a**, Release from **POE-C9G1**, demonstrating how the inclusion of hydrophobic (3% (w/w) S65) or basic (5% (w/w) PEI) additives can shift the release window from one month (absence of additives) to a projected duration exceeding 12 months while retaining zero-order profiles. **b**, Release curves from **POE-C9T1** with Mg salts (MgSt or Mg(OH)_2_ nanoparticles, 3% (w/w)), further supporting the role of basic additives in slowing down release rates. **c**, Comparative release curves obtained from **POE-H3C6P1** showcasing sustained zero-order release, and commercial PLGA (Resomer^®^ RG 756 S) displaying irregular release rate with significant delayed burst. **d**, Release curves obtained from **POE-H3C6P1** loaded with 5% (same data from **Fig. 3c**) and 10% (w/w) of LNG, demonstrating equivalent cumulative release, and a sustained zero-order release over 300 days. **e**, Representative Raman confocal maps showing the spatial distribution of LNG and the lack of water in **POE-H3C6P1** MPs loaded with 5% (w/w) LNG. **f**, Raman confocal maps showing the spatial distribution of LNG and swelling behavior of PLGA MPs (Resomer^®^ RG 503 H) loaded with 5% (w/w) LNG. Scale bars: 30 µm.

MPs fabricated from POEs containing hydrophilic diols (**POE-C9T1, POE-H3C6T1, POE-C9G1**) showed a “fast” release profile, with the full LNG dose released over 1 to 2 months (**Fig. 3a-b** and **Extended Data Fig. 1**). To prolong LNG release, we included in the MPs either hydrophobic additives, i.e. Span^®^ 65 (S65), or basic additives, i.e. polyethylenimine (PEI), magnesium hydroxide nanoparticles (Mg(OH)_2_ NPs) or magnesium stearate (MgSt). Previous studies on macroscopic POE implants have shown that basic additives can neutralize acidic degradation products, thereby preventing acid-induced auto-catalytic degradation of polymer matrices, eventually slowing down drug release rates^46, 47^. Building on this, we hypothesized that incorporating such additives into our system could enhance stability against acid-induced auto-catalytic degradation of the polymer matrix. The inclusion of the selected additives in the MP formulation enabled us to extend the release window and control its duration for both **POE-C9G1** and **POE-C9T1** (**Fig. 3a-b**).

We then tested MPs fabricated with POEs that included only hydrophobic diols. **POE-H1C2** showed complete drug release over a 7-month period, albeit displaying an initial lag time of approximately 1 month. Incorporation of Mg(OH)_2_ NPs or MgSt during the MP fabrication process prolonged the release window, consistent with the effect observed for the more hydrophilic polymers. A similar effect was seen for MPs produced from **POE-D1C2**, displaying a release window within a 12-month period. However, releases observed from both **POE-H1C2** and **POE-D1C2** displayed irregular profiles with extended lags both at the beginning and throughout the release window (**Extended data Fig. 2**).

Based on these findings, we envisioned that the addition of tertiary amine-containing piperazine diol **P** to POE formulations could result in a polymer structure with Brønsted basic functionality to prolong zero-order LNG release from POE MPs without additives. Furthermore, we included a hydrophilic diol to avoid multistage release behaviors as observed when utilizing only hydrophobic diols (i.e. **POE-H1C2** and **POE-D1C2**). In this regard, we modified POE-H1C2 with the inclusion of monomer **P**, leading to the polymer matrix **POE-H3C6P1**.

**POE-H3C6P1** demonstrated sustained zero-order kinetics, with ∼40% of the total dose released after 6 months (**Fig. 3c**). By contrast, MPs fabricated from a commercially available slow-degrading PLGA polymer^48^ (Resomer^®^ RG 756 S, 75:25 ratio of L/G, *M*_w_= 76-115 kDa) displayed an irregular release profile characterized by a significant delayed burst-release, resulting in a sudden accelerated release of LNG after 120 days, as commonly observed with PLGA-based formulations, indicating their limited suitability for applications requiring highly controlled, long-term release (**Fig 3c**). Increasing the loading of LNG in **POE-H3C6P1** from 5% (w/w) to 10% (w/w) did not cause any burst release, with MPs prepared with this polymer displaying cumulative release independent of the drug loading. **POE-H3C6P1** with 10% (w/w) LNG loading showcased a consistent linear release, reaching ∼60% of the initial dose in ∼300 days *in vitro* (**Fig. 3d**).

To further elucidate the mechanism of release from **POE-H3C6P1**, we performed Raman confocal imaging longitudinally over a period of 58 days (**Fig. 3e** and **Supplementary Fig. 3-4**). This technique enables chemical mapping of individual MPs, allowing assessment of water ingress into the particles and spatial distribution of LNG within the particles. MPs loaded with both 5% (w/w) and 10% (w/w) LNG showed a stable LNG signal across the surface of the particles, with no water ingress over time, consistent with a surface erosion-mediated release. By contrast, Raman images of PLGA MPs (Resomer^®^ RG 503 H, 50:50 ratio of L/G, *M*_w_= 24-38 kDa) exhibited significant swelling and a drastic drop in LNG signal across the whole particle surface over time, consistent with a bulk release mechanism (**Fig. 3f**).

Following the *in vitro* release studies, varying concentrations (1, 5 and 50 mg/mL) of freshly prepared and partially degraded **POE-H3C6P1** MPs were assessed for their potential acute cytotoxic effect on dermal fibroblast cells. Partially degraded MPs were incubated at 37 °C for one month prior to being introduced to the cell culture to simulate the effect of degradation. An AlamarBlue assay was employed to assess cell viability of dermal fibroblasts. No cytotoxicity was observed upon exposure to freshly prepared MPs or degraded MPs (**Supplementary Fig. 5**). These results indicate good biocompatibility of the **POE-H3C6P1** formulation consistent with previously reported POE formulations^49, 50^.

### Microparticles-hydrogel composite shows *in vivo* injectability and retrievability

To evaluate the ease of injectability of MP/HG composites through 23G needles, we developed a custom setup using a Bose^®^ 3200 Test Instrument to quantify the injection force required for the administration of the LNG-free composite, following established protocols^51, 52^. For this, the syringe plunger was compressed by the crosshead at a fixed rate, and the resulting force was recorded. We tested an unloaded HG, a high particle concentration MP/HG composite formulation (19.3% (w/v)), and its corresponding reconstituted formulation after freeze-drying. All samples were injectable through a 23G needle, with an average injection force below 20 N (**Fig. 4a**), suitable for subcutaneous administration^53^.

**Fig. 4:**
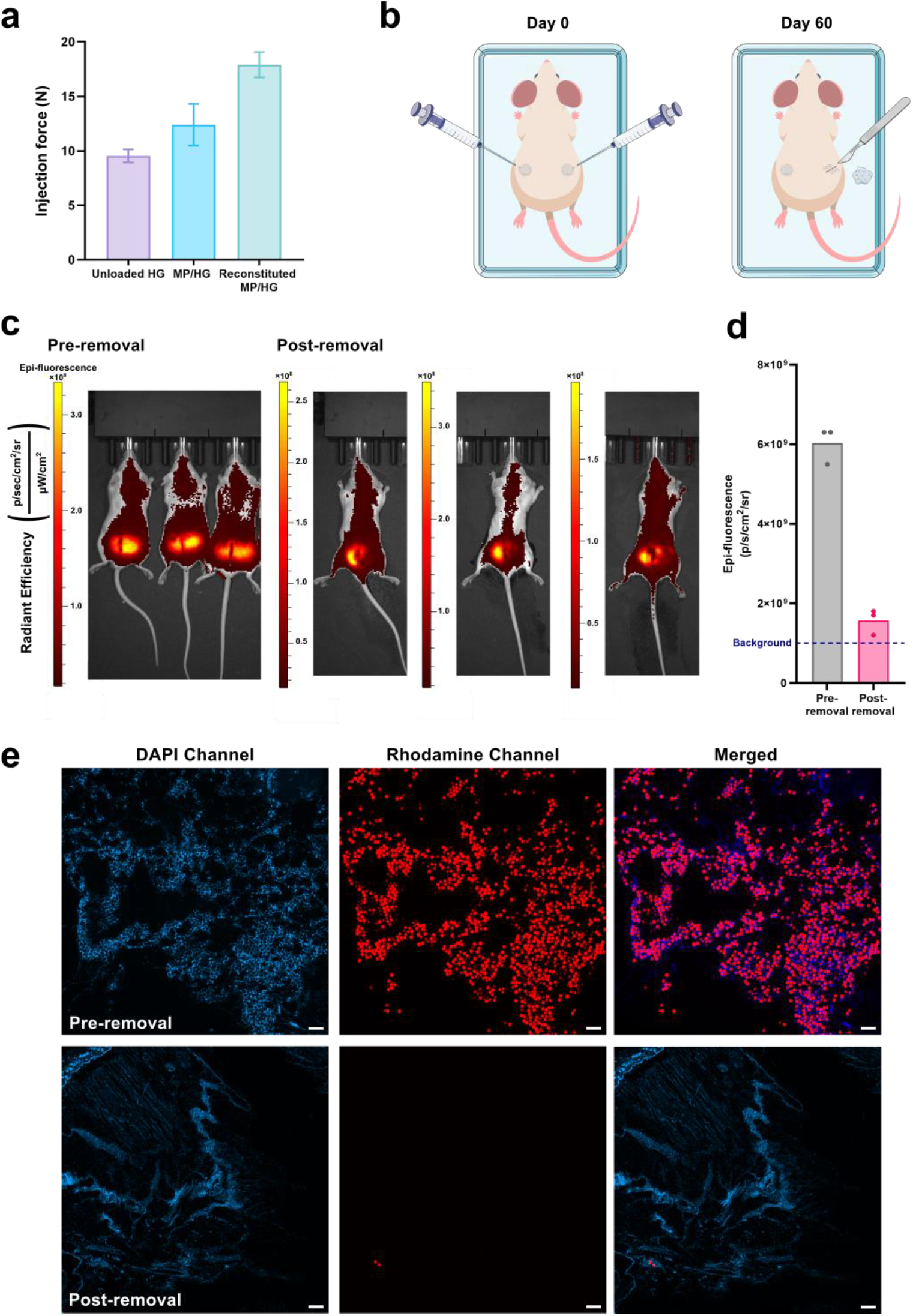
*In vivo* injectability and retrievability of the MP/HG composite. **a**, Quantitative evaluation of the injection force through a 23G needle of particle-free, particle-loaded and reconstituted HG (Bose^®^ test instrument). Data represents mean ± s.d. (n = 3). **b**, Schematic of the injection and removal procedure 60 days post injection. **c**, IVIS images of mice (n = 3) at day 60 pre- and post-removal of MP/HG composite. **d**, IVIS signal intensity comparison pre- and post-removal of the MP/HG implant by incision. **e**, Representative images of tissue sections analyzed by fluorescence microscopy. Scale bar: 300 μm.

To assess the stability of the implant and its safe removal, 150 µL of Cy7-Rhodamine B covalently co-labeled PLGA MP/HG composite (19.3% (w/v)) was subcutaneously injected in the left and right sides of the back of female BALB/c mice through a 23G needle (**Fig. 4b**). Mice were imaged *via* IVIS^®^ on the day of the injection, and a strong Cy7 signal post-implantation confirmed the successful injection of the composite. For each mouse, the implant on the right side was removed at day 60 through a small incision and gentle squeezing at the injection site, and the mice were imaged after removing the composite (**Fig 4c**). We compared the total signal at the injection site pre- and post-removal of the implant, and a significant reduction (*ca*. 90%) in the Cy7 signal was observed, indicating removal of MPs *via* hydrogel explantation (**Fig. 4d**). Tissue sections were further investigated by fluorescence microscopy, showing how the removal of the composite by incision provided substantial removal of the MPs at 60-day post-implantation (minimal number of particles detected in N = 15 images, indicative example in **Fig. 4e**, further images provided in **Supplementary Fig. 6**). Images of the extracted gels show that the majority of the particles stayed within the gel after removal (**Supplementary Fig. 7**).

### *In vivo* pharmacokinetic studies in rats confirm sustained zero-order release of LNG

Following encouraging observations of zero-order *in vitro* release, we assessed the pharmacokinetic (PK) behavior of LNG in a rat model. To evaluate LNG PK from POE MP/HG composites, female Sprague-Dawley rats were administered our best-performing candidate for long-term release **POE-H3C6P1** with different LNG loadings (either 5 or 10% (w/w)), and levels of LNG in plasma were measured over 28 days. Rats were injected with composites containing a total amount of 9 mg of LNG (**Supplementary Table 1**). Rats that received LNG-loaded POE MP/HG composites exhibited stable LNG plasma concentrations over the 4-week period, consistent with the linear release observed *in vitro*. Furthermore, we observed no significant difference in plasma LNG concentrations between the 5% and 10% (w/w) LNG-loaded **POE-H3C6P1** MPs groups (**Extended Data Fig. 3**). This loading-independent behavior aligns with the hypothesized surface-erosion mechanism, where the release of the drug is independent from the loading per particle, but depends only on the total amount of drug administered.

Following this initial study, we performed a second study over 161 days, comparing MP/HG composites prepared with 10% (w/w) LNG-loaded **POE-H3C6P1** or 5% (w/w) LNG-loaded **POE-C9T1/MgSt**, and a total dose of ∼9 mg of LNG (**Supplementary Table 2**), equivalent to the clinically used dose of 30 µg/day over 300 days. Here we sought to compare two different formulations, both displaying a linear release profile *in vitro*, albeit with very different release rates, with the aim of demonstrating the adaptability of the platform for various time scales and potentially different applications. In this study, we also compared our MPs formulation with a hydrogel loaded with LNG crystals, as these have been employed in long-acting LNG formulations^11^ which rapidly peaks at day one and then slowly decays (**Fig. 5a**). Assessing the use of the faster releasing **POE-C9T1/MgSt** MP/HG composites as an implantable contraceptive solution, we observed a stable concentration profile over ∼49 days, followed by a decay over ∼28 days (**Fig. 5b**). Rats administered with 10% (w/w) LNG-loaded **POE-H3C6P1** MP/HG composites aiming for a longer efficacy window showcased an extremely stable plasma concentration profile over the course of the study (**Fig. 5c**), consistent with the zero-order release rate observed *in vitro*. Furthermore, the profiles measured in both PK studies (using different batches of particles produced with different batches of polymer) exhibited substantial overlap, supporting the robustness of our platform (**Supplementary Fig. 8**). Both POE formulations displayed a prolonged and steadier release profile compared to the LNG crystals control.

**Fig. 5:**
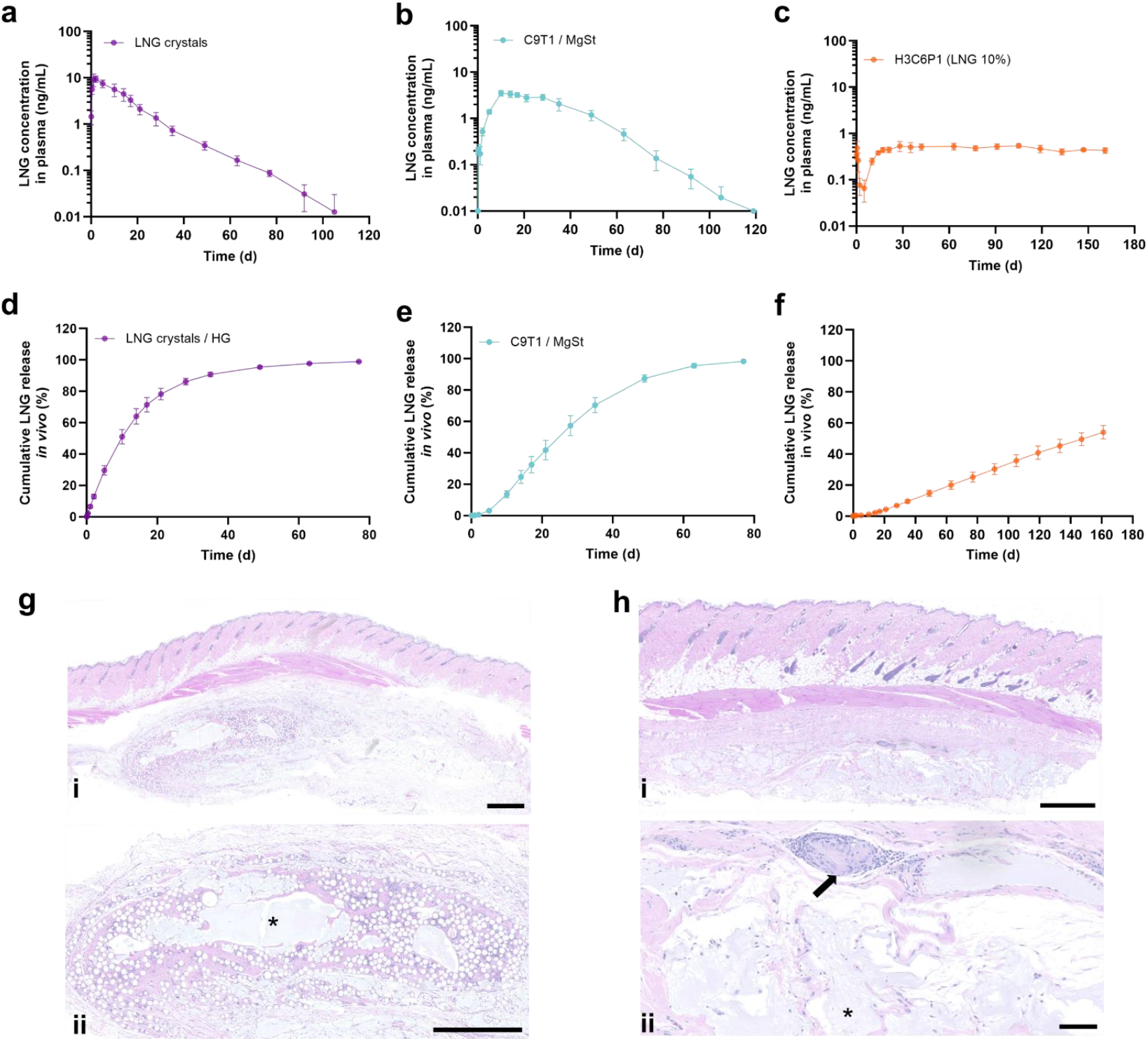
Pharmacokinetics, physiological effects, and histology of LNG in rats. **a-c**, Plasma LNG concentration over time following administration of three different formulations, (n = 5). **d– f**, Cumulative *in vivo* LNG release over time in three corresponding treatment groups. The plasma detection limit for LNG was 0.025 ng·mL^−1^. LNG absorption was estimated using the Wagner–Nelson method, with numerical deconvolution applied to concentration profiles. Data represent mean ± s.d. (n = 5). (**a**,**d**, LNG crystals in HG; **b**,**e**, 5% (w/w) LNG-loaded **POE-C9T1/MgSt** MP/HG; **c**,**f**, 10% (w/w) LNG-loaded **POE-H3C6P1** MP/HG composite). **g**, Representative H&E-stained tissue sections collected at the end of the study (day 168) from animals which received 10% (w/w) LNG-loaded **POE-H3C6P1** MP/HG composite. *i*, the overlying epidermis is intact, and the fascia beneath the panniculus carnosus is expanded by a nodule composed of mild fibrous connective tissue with rare multinucleated macrophages (Langhans cells) and minimal lymphocytic infiltration. *ii*, No capsule demarcates the nodule, which blends with the surrounding fascia; a moderate amount of pale extracellular material (asterisk), consisting of the gel, separates the fibroblasts. Scale bar: 1 mm. **h**, Tissue sections from injection sites treated with 5% (w/w) LNG-loaded **POE-C9T1/MgSt** MP/HG composite. *i*, The overlying epidermis is intact, and the fascia beneath the panniculus carnosus is expanded by a poorly defined nodule. *ii*, The nodule consists of mild fibrous connective tissue with rare multinucleated macrophages (Langhans cells; arrow) and minimal lymphocytic infiltration. Inflammation is minimal, no capsule is present, and moderate amounts of pale basophilic extracellular material (asterisk), consisting of the implant, are observed. Scale bar: 100 µm.

The LNG plasma concentration profiles obtained from the composites were then converted into cumulative release curves using the Wagner-Nelson method^28^ and compared with the cumulative releases obtained *in vitro* (**Fig. 5d-f**). The curves obtained from the *in vivo* study for the two **POE** formulations show a zero-order profile with no observed burst, consistent with the *in vitro* results, albeit the *in vivo* data displayed a faster rate, consistent with other studies on polymeric drug release implants.^28^ Comparing the curves obtained from **POE-H3C6P1** and **POE-C9T1/MgSt**, the former slow-release formulation achieved ∼58% release of the total dose compared to the latter, fast-release one (**Fig. 5b-c** and **Supplementary Fig. 9**), indicating the potential for the slow degrading platform to possess desirable release behavior for up to 12 months.

Microscopic examination of the injection site 168 days post-injection revealed samples with no overt evidence of inflammation or with minimal inflammatory infiltrates, with minimal fibrosis and without sign of capsule formation **(Fig. 5 h-i)**. The epidermis showed no evidence of erosion, ulceration or hyperplasia, indicating rapid healing after injection of the implant. Overall, these results support the tolerability of our soft implant.

## Discussion

In this study we demonstrate the potential of POE MP/HG composites as a long-acting, minimally invasive drug release platform. By utilizing bespoke surface-eroding POE formulations, this system enables sustained, zero-order release of LNG with a tunable release profile from 1 month to a projected release of over 12 months *in vitro* depending on the POE structure. Our soft implant displays a zero-order release profile *in vivo*, with an unprecedented stable plasma concentration over the targeted release window. Depending on the patient’s need, we could exchange the POE formulation and control the length of the release window, while retaining the zero-order release rate. The injectable nature of the composite ensures ease of administration with a clinically relevant 23G needle, reducing barriers associated with surgical implantation required for many existing long-acting formulations. Additionally, the hydrogel carrier enables retrievability even after prolonged implantation, providing an effectively reversible long-acting formulation. This study represents a significant advancement in developing next-generation, easy-to-administer long-acting formulations that balance long-term efficacy with reversibility and convenience for clinicians and patients. As the drug release rate of our soft implant can be effectively controlled through polymer degradation, we envision this system used for different therapies beyond contraception, where repeated dosing over an extended period of time is required, including diabetes, infectious diseases, chronic inflammations, and autoimmune diseases.

## Methods

### Materials

All chemicals were purchased from Merck and used without further purification unless otherwise specified. Medical grade HA fillers (Belotero Intense) were purchased from Fillerworld. All other solvents and chemicals were of analytical grade and stored at room temperature.

### Synthesis of DETOSU

3,9-Diethylidene-2,4,8,10-tetraoxaspiro[5.5]undecane (DETOSU) was prepared through the isomerization of 3,9-divinyl-2,4,8,10-tetraoxaspiro[5.5]undecane (DVTOSU) according to a previously described method^54^ (**Supplementary Scheme 1)**. Briefly, potassium *t*-butoxide (10.8 g, 96.24 mmol) and 50 mL of ethylenediamine were mixed under a nitrogen atmosphere with a reflux condenser. After allowing this mixture to stir for 15 min, DVTOSU (9 g, 42.4 mmol) was added to the flask, and the mixture was heated to 100 °C under nitrogen atmosphere for 16 h. The next day, the mixture was poured into 600 mL of cold water, and the product was extracted with pentane and dried with K_2_CO_3_. The crude product was purified by distillation over CaH_2_ in a short-path distillation apparatus (pressure ∼3 mbar, bath temperature 125 °C - 145 °C) to isolate DETOSU. Yield was calculated as 25% after distillation. The ^1^H NMR resonances (**Supplementary Fig. 10**) and a characteristic IR band of DETOSU at 1700 cm^−1^ (**Supplementary Fig. 11**) were used to further confirm the synthesis of the desired molecule. FTIR spectra of the synthesized DETOSU were identical to those reported in the literature.^55^

### Synthesis of POEs

All POEs were synthesized through step-growth polymerization of DETOSU with a diol or a mixture of diols. Under strictly anhydrous conditions, a solution of various diols (1,6-hexanediol (H), 1,10-decanediol (D), *trans-* 1,4-cyclohexanedimethanol (C), triethylene glycol (T), glycerol monomethacrylate (G), and 1,4-bis(2-hydroxyethyl)piperazine (P)) at pre-defined ratios (**Table 1**) in 6 mL anhydrous THF and DETOSU (50 mol% relative to the total amount of diols) in 4 mL THF were mixed under nitrogen atmosphere. The reaction was catalyzed by 50-100 µL of iodine/pyridine mixture (500 mg/mL, addition rate of 1 drop every 10 seconds) and carried out at room temperature. An exothermic reaction was observed, indicating the initiation of polymerization. Polymerization was terminated after 1 hour, and the polymers were precipitated into cold methanol with 5 drops of NEt_3_. The polymers were then dried under vacuum. POEs were obtained with quantitative yields, and all the structures were confirmed by ^1^H-NMR (**Supplementary Fig. 12-18**).

### Fabrication of MPs

POE and PLGA microparticles were fabricated using a custom-made microfluidic flow focusing set-up, consisting of two coaxially aligned glass capillaries, similar to the set-up reported by Vladislavjevic et al.^42^. The outer glass capillary has an inner diameter (I.D.) of 1.4 mm, while the inner capillary has an I.D. of 210 µm. The continuous phase was an aqueous solution (Millipore, deionized) of phosphate buffer (PB, pH = 7.4, 10 mM) containing Mowiol^®^ 4-88 (1% (w/v), *M*_w_= ∼31 kDa) as a surfactant. The disperse phase was dichloromethane (DCM) containing polymer (75 mg/mL). Digitally controlled syringe pumps delivered both liquid phases to the microfluidic device at constant flow rates. Flow rates of 50 µL/min for the dispersed phase and 150 µL/min for the continuous phase were used to obtain particles with diameters of 80 µm. PTFE tubing connected the syringes with the inlets of the device and the outlet of the device with a glass petri dish containing 13 mL of PVA solution in PB buffer (1% (w/v)). DCM within the droplets was removed by evaporation overnight. After removal of DCM, solidified microparticles were collected in a 50 mL conical tube and washed with deionized water (30 mL) 5 times to remove excess PVA. The particle suspension was then frozen and lyophilized under reduced pressure to remove the continuous aqueous phase. PLGA microparticles were produced with the same procedure. For the *in vivo* injectability and retrievability study, the dispersed phase consisted of Resomer^®^ RG 756 S (75:25 ratio of L/G, *M*_w_= 76-115 kDa) at 72 mg/mL, Cy7 PLGA (see next section for details on the polymer) at 1.5 mg/mL and PLGA-Rhodamine B (50:50 L/G, 20 kDa, Nanosoft Polymers™) at 1.5 mg/mL.

### Cy7 labeling of PLGA for *in vivo* imaging

The synthesis followed established protocols.^56^ Briefly, Resomer^®^ RG 504H (poly(D,L-lactide-*co*-glycolide, acid terminated, L/G: 50:50, Mw= 38-54 kDa) was dissolved in DCM at 0.01 mmol/mL. To this, 5.6 equivalents of *N,N′*-dicyclohexylcarbodiimide and *N*-hydroxysuccinimide, each pre-dissolved in DCM at 0.5 mmol/mL, were added. The reaction was stirred at room temperature until precipitation occurred, then continued for an additional hour. The solution was filtered through a cotton-packed pipette to remove dicyclohexylurea. In a separate vial, Cy7-amine was dissolved in DMF at 0.027 mmol/mL and added to the PLGA–NHS mixture. Also, 4 equivalents of triethylamine, diluted in DMF at 0.13 mmol/mL, were added. The conjugation was maintained under magnetic stirring for 18 h at room temperature. The mixture was then precipitated into ice-cold methanol at a 1:10 volume ratio and re-precipitated in cold methanol three times. Finally, the product was dialyzed using a Pur-A-Lyzer™ Mini Dialysis Kit against DCM for 3 days to remove the unreacted dye, protected from light with aluminum foil. After dialysis, the precipitate was recovered in ice-cold methanol and dried in a vacuum oven for 2 days.

### Synthesis of Mg(OH)_2_ nanoparticles

Mg(OH)_2_ synthesis was adapted from Lih et al.^57^ 20 mL of an aqueous Mg(NO)_3_ solution (130 mg/mL) were added dropwise to 20 mL of an aqueous solution of NaOH (36 mg/mL) under stirring. The solution was further stirred for 30 min, before the product of the precipitation reaction was centrifuged and oven dried at 60 °C overnight. The obtained product was then coated with oleic acid by dissolving the dried powder at 5 mg/mL in a H_2_O:EtOH solvent 1:1 (v/v) together with 5 mg/mL of oleic acid. The solution was kept stirring overnight, then the product was centrifuged and washed with DCM 5 times before use. TEM images of the nanoparticles were taken with a JEM-2100plus (JEOL) and are provided in **Supplementary Fig. 19**.

### LNG release from POE MPs *in vitro*

To evaluate the *in vitro* release of LNG from MPs, we used the release medium of PBS (137 mM NaCl, 2.68 mM KCl, 10.14 mM Na_2_HPO_4_, 1.76 mM KH_2_PO_4_) containing 3% (w/v) 2-hydroxypropyl-*β*-cyclodextrin and 0.1% (w/v) sodium azide. Approximately 5 mg of POE microparticles were placed into 30 mL of release medium in glass vials together with ∼25 µL of hydrogel. The vials were incubated in a shaker at 37 °C and were shaken at 100 rpm. At predetermined time points (every 7 days), 100 µL of release medium was collected and replaced with the same amount of fresh medium. Collected samples were analyzed by reversed-phase HPLC (Agilent 1260 Infinity) equipped with a UV detector to quantify the LNG concentration.

### Preparation of MP/HG composites

MPs were weighed into an Eppendorf tube and incorporated into the HG at the desired concentration through vortexing for 5 min, followed by manual mixing with a spatula. The resulting MP/HG mixture was then freeze-dried and stored at –20 °C.

### Reconstitution process of MP/HG composites

The freeze-dried MP/HG composites were re-suspended in Milli-Q water to achieve the initial hydrogel weight. The samples were then incubated at room temperature on a Thermomixer set to 350 rpm for 24 h, then stored at 4 °C.

### Injection Force Measurement Using Bose^®^ 3200 Test Instrument

Injection force measurements were performed using a Bose ElectroForce 3200 testing instrument equipped with a custom-built setup for quantitative injection force acquisition. The setup comprised a stainless-steel support and a 3D-printed syringe holder to ensure consistent positioning and alignment during testing. A crosslinked, medical-grade hyaluronic acid (HA) filler was tested in three formulations: particle-free, HA hydrogels loaded with PLGA microparticles (19.3 wt%) and re-constituted formulation. Each formulation was extruded through single-use syringes fitted with 23G needles. Injections were conducted at a constant crosshead speed of 0.4 mm s^−1^ over a total displacement of 4 mm.

### Rheological Characterization of the MP/HG Implant

Rheological measurements were performed on particle-free and HA hydrogels loaded with PLGA microparticles (6.4% and 19.3% (w/w)) using an AntonPaar compact rheometer (MCR203) at 37 °C. Frequency sweeps (0.1% strain, 0.1-10 Hz) were conducted to assess viscoelastic properties. Storage modulus (G′) and loss modulus (G″) were evaluated at 0.7 Hz to compare formulations.

### Mouse Strains

Breeding and all animal procedures were performed in accordance with UK Home Office regulations under project license PP5920580. Female mice (8-10 weeks old) were used in all experiments. Balb/c mice were provided by The Francis Crick Biological Research Facility.

### *In vivo* injection and retrieval of MP/HG composites

150 µL of Cy7-Rhodamine B co-labeled PLGA MP/HG composite (19.3% (w/v)) was loaded in syringes equipped with 23G needles. Animals were anaesthetized with isoflurane and gels were injected subcutaneously on each flank. Mice were euthanized using cervical dislocation on day 60 and a small incision was performed using a scalpel to remove the hydrogel. After fluorescence imaging, the surrounding tissue was collected and fixed with 4% paraformaldehyde overnight at 4 °C, washed 3 times with PBS, embedded in cryomoulds using OCT embedding medium and stored at -80 °C. Blocks were then sectioned for histology into 10 μm sections using the cryostat (Leica, CM3050S).

### Immunostaining

For immunostaining, sections were thawed at room temperature for 10 min and stained with DAPI at 1:500 dilution for 45 min at room temperature in the dark. Slides were washed with PBS 3 times for 5 min. Finally, slides were rinsed in distilled water and mounted using Vectashield Vibrance^®^ antifade mounting medium (Vector Laboratories) and #1 Menzel coverslips (Thermo Fisher Scientific).

### *In vivo* Imaging

*In vivo* fluorescence imaging was done using the IVIS^®^ molecular imaging platform (Perkin Elmer) measuring the radiant efficiency in units of (photons/sec/cm^2^/sr)/(μW/cm^2^) for Cy7 signal. Mice were imaged on day 60 before and after hydrogel removal.

### Cytocompatibility Assessment of POE-H3C6P1 MPs

Three batches of **POE-H3C6P1** MPs were separately weighed in Eppendorf tubes and sterilized under UV light for 30 min and medium was added to achieve a slurry of particles of 200 mg/mL. Dermal fibroblast cells were seeded at 20×10^3^ cells/well in a 96-well plate overnight at 37 °C. For degraded particles, MPs (200 mg/mL) were incubated in medium at 37 °C for 30 days. Freshly prepared MPs (three technical replicates per batch for each condition) or degraded MPs (four technical replicates per batch for each condition) were added to the wells at 1, 5, or 50 mg/mL and incubated for 24 h at 37 °C. Treatment media was removed and replaced with 10% (v/v) AlamarBlue. After 2 h of further incubation, cell viability was determined by reading fluorescence at 570 nm excitation/585 nm emission. Relative cytotoxicity and viability were calculated relative to buffer-only controls.

### High Performance Liquid Chromatography (HPLC)

Reversed-phase HPLC–UV analysis was performed using an Agilent 1260 Infinity Quaternary LC equipped with an autosampler (G1329A), 1260 Infinity II Diode Array Detector (G7115A). Measurements were run using a Gemini Phenomenex NX-C18, 150 × 4.6 mm, particle size = 5 μm, pore size = 110 Å, at a flow rate of 1 mL/min and injection volume = 50 μL. The column was equipped with a SecurityGuard ultra cartridge (particle size: 5 μm), Phenomenex. Instrumental control of the HPLC system as well as data collection, processing and integration used ChemStation software.

### Optimized HPLC method for LNG detection

A standard LNG stock solution (60 µg/mL) was prepared in release medium of PBS containing 3% (w/v) 2-hydroxypropyl-β-cyclodextrin and 0.1% (w/v) sodium azide at 37 °C. LNG calibration standards: 40, 20, 10, 5, 1, 0.5, and 0.1 µg/mL were then prepared by serial dilution in release buffer to create a calibration curve, 0.1 to 40 µg/mL. The optimal chromatographic conditions for LNG detection consisted of a gradient elution of HPLC grade acetonitrile with 0.1% (v/v) formic acid and HPLC grade water with 0.1% (v/v) formic acid, at a flow rate of 1 mL/min, starting at 50:50 (v/v), changing to 95:5 (v/v) at 8 min and then returning to 50:50 (v/v) at 9 min. The injected sample volume was 50 µL, and LNG absorbance was recorded at 254 nm. The column temperature was set to 40 °C, and a single run lasted 13 min.

### Gel Permeation Chromatography (GPC)

Molecular weight characteristics of all POEs were characterized using an Agilent PL GPC-50 instrument equipped with a refractive index (RI) detector running in GPC grade DMF (containing 0.075% (w/v) LiBr) at a flow rate of 1.0 mL/min at 40 °C through two GRAM Linear columns (Polymer Standards Service) in series. Near monodisperse poly(methyl methacrylate) standards dissolved in eluent were used to calibrate the instrument. Polymer was dissolved in eluent at 5 mg/mL and filtered through a 0.2 μm syringe filters prior to analysis.

### Nuclear Magnetic Resonance Spectroscopy (NMR)

^1^H and ^13^C NMR spectra were acquired on a Bruker Avance 500 MHz NMR spectrometer operating at 293 K. Deuterated chloroform (CDCl_3_) was acquired from Sigma Aldrich and used as received. Chemical shifts (δ) were referenced to the residual solvent peak (δ = 7.26 ppm). Proton (^1^H) NMR data are reported as chemical shifts with the following multiplicity notation: s = singlet; d = doublet; t = triplet; q = quartet; m = multiplet; br = broad. This is followed by the proton position and then the coupling constants (*J*) in Hertz if applicable.

### Fourier Transform Infrared Spectroscopy (FTIR)

AT-FTIR analysis on a Perkin Elmer Spectrum 100 spectrometer at a resolution of 1 cm^-1^ with 48 scans were performed for the analysis of synthesized DETOSU.

### Scanning Electron Microscopy (SEM)

Microparticles were transferred to conductive tape on the top surface of a metal stub. Cross-sections were prepared by mixing particles with OCT embedding medium in cryomoulds and stored at -80 °C. Blocks were then sectioned using a cryostat (Leica, CM3050S) to reveal particles in cross-section, and transferred to conductive tape. All samples were coated with gold (30 seconds deposition time, 20 mA current) *via* sputtering deposition (Emitech K575X peltier cooled) before scanning electron imaging was conducted on a Zeiss Leo Gemini 1525 microscope in secondary electron mode, at 3 kV accelerating voltage, 10 mm working distance, and 30 µm aperture.

### Confocal Raman Imaging Microscopes

MPs were imaged with a WITec alpha 300R + Raman confocal microspectrometer equipped with a piezoelectric stage (UHTS 300, WITec GmbH), 63X water immersive objective lens (Zeiss W Plan Apochromat 63X, N.A = 1), red solid-state excitation laser (λ = 785 nm, 85 mW, WITec GmbH) and an imaging spectrograph (Newton, Andor Technology Ltd). This setup enabled acquisition of spectral data across a wavenumber range up to 3600 cm^−1^.

### *In vivo* pharmacokinetic study

The PK study was performed by Sygnature Discovery in female Sprague Dawley rats (281.2 g ± 10.5 g). All procedures involving animals were carried out in accordance with the Animals Scientific Procedures Act 1986/ASPA Amendment Regulations 2012 and under the authority of UK Home Office Project License PPL0194953. Plasma samples were collected at predetermined intervals following subcutaneous (SC) administration of the different formulations tested. The amount of levonorgestrel in each sample was quantified by LC-MS/MS against a calibration curve. The Wagner-Nelson method was employed to generate cumulative release profiles. Since formulation H3C6P1 only released part of the total dose over the duration of the experiment, the cumulative release for H3C6P1 was obtained normalizing against the average final AUC of the other two formulations.

## Data Availability

Raw research data is available upon request from the corresponding author.

## Acknowledgements

This work was supported in part by the Gates Foundation (INV004356). The findings and conclusions contained within are those of the authors and do not necessarily reflect positions or policies of the Gates Foundation.

Under the grant conditions of the Foundation, a Creative Commons Attribution 4.0 License has already been assigned to the Author Accepted Manuscript version that might arise from this submission. L.P., D.J.P. and J.Y. were supported by the European Union’s Horizon 2020 research and innovation programme under Marie Skłodowska-Curie Actions (101106805, 101027174, 839137). L.P. acknowledges support from the CRUK early detection and diagnosis primer award (100017). A.N. acknowledges support from a Sir Henry Wellcome Postdoctoral Fellowship (209121_Z_17_Z) from the Wellcome Trust. T.R. and I.M. were supported by the Francis Crick Institute, which receives its core funding from Cancer Research UK (CC2051), the UK Medical Research Council (CC2051), the Wellcome Trust (CC2051), and the European Research Council grant (ERC CoG-H2020725492). R.X., R.B. and M.M.S. acknowledge funding from the Engineering and Physical Sciences Research Council (EP/T020792/1). Y.C. gratefully acknowledges support from the President’s PhD Scholarship at Imperial College London. M.M.S. acknowledges support from the Department for Science, Innovation and Technology and the Royal Academy of Engineering Chair in Emerging Technologies award (CiET2021\94), the University of Oxford Strategic Research Fund and the Rosetrees Trust. We kindly acknowledge Jonathan P. Wojciechowski for HPLC method development for the *in vitro* drug release studies. Please note that works submitted as preprints have not undergone peer review.

## Author Contributions

C.K. and L.P. contributed equally to this work. C.K. was responsible for the design, synthesis and characterization of polymer matrices, optimizing the fabrication of drug-loaded polymeric microspheres, aiding *in vitro* drug release analysis and aiding in encapsulation microspheres into hydrogel for *in vivo* analysis. L.P. was responsible for the fabrication of drug-loaded polymeric microspheres, and for designing and overseeing the *in vivo* PK study. A.N. was responsible for performing the *in vitro* drug release analysis. T.R. was responsible for carrying out *in vivo* mouse experiments. A.S. contributed to polymer design and was responsible for aiding the synthesis and characterization of polymer matrices. G.B. was responsible for optimizing the encapsulation of microspheres into hydrogel. D.J.P. contributed to polymer design, established *in vitro* drug release analysis, and contributed to *in vivo* study design. L.Z., T.F.D. and M.C. were responsible for SEM imaging. P.P. was responsible for Raman confocal imaging. R.X. was responsible for *in vitro* cell biocompatibility assays. A.P.C. aided in the *in vitro* drug release studies. Y.C. performed TEM imaging. J.Y. and F.S. supported the design of the POE library. A.S-B. has performed histopathological examination and interpretation of tissue samples. I.M. supported the *in vivo* retrievability study. C.K. and L.P. drafted the paper with input from all co-authors. M.M.S. supervised the study.

## Extended Data

**Extended Data Fig. 1:**
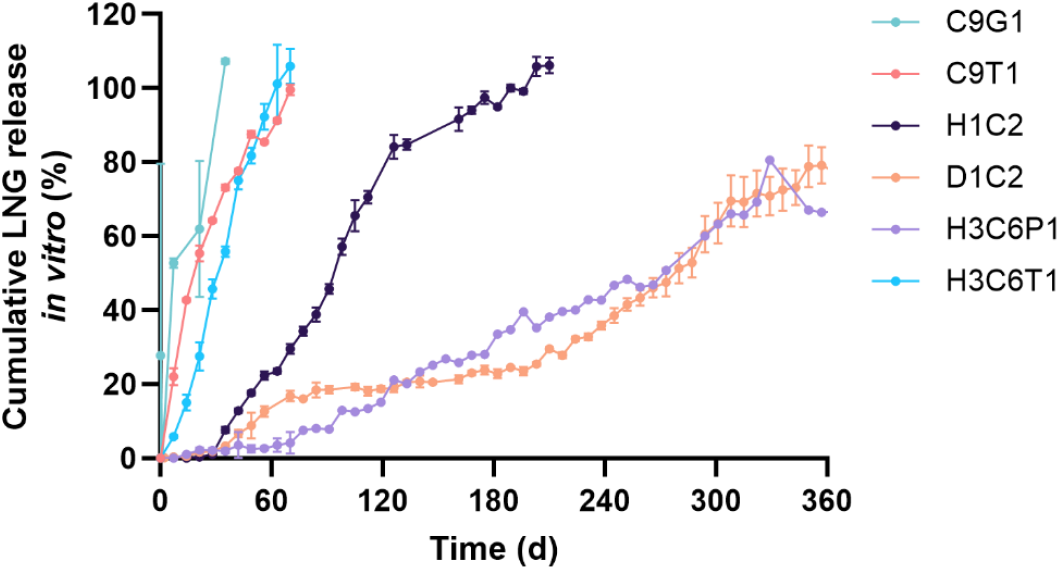
Cumulative *in vitro* release of LNG from MPs formed by different POE formulations containing 5% (w/w) LNG MPs incubated in PBS containing 3% (w/v) 2-hydroxypropyl-β-cyclodextrin and 0.1% (w/v) sodium azide at 37 °C, plotted over time. Data represent mean ± s.d. (n = 3). A subset of this data is also shown in **Fig. 3**.

**Extended Data Fig. 2:**
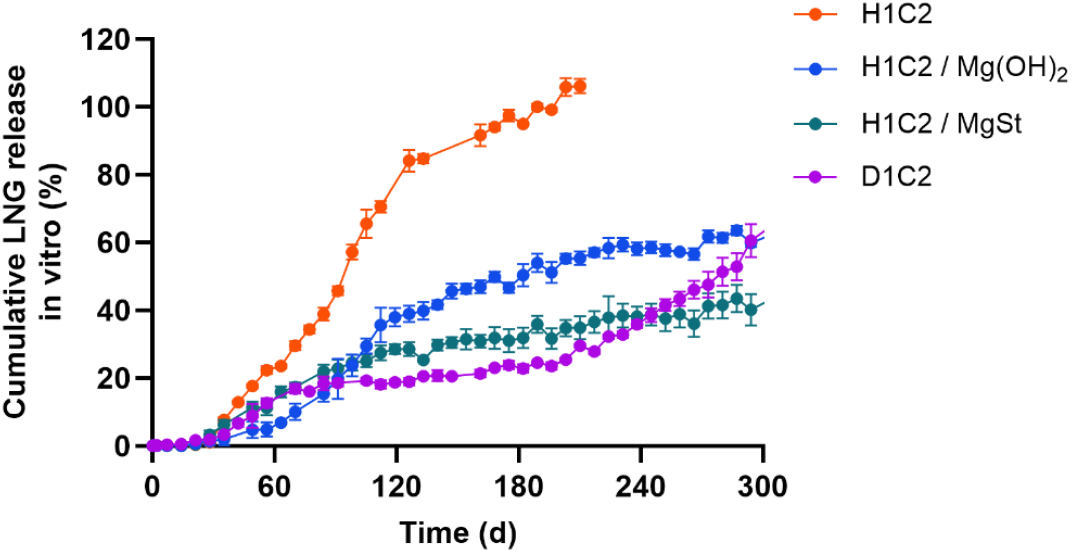
Cumulative *in vitro* release of LNG from POE-H1C2 and POE-D1C2 MPs containing 5% (w/w) LNG MPs incubated in PBS containing 3% (w/v) 2-hydroxypropyl-β-cyclodextrin and 0.1% (w/v) sodium azide at 37 °C, plotted over time. Data represent mean ± s.d. (n = 3). A subset of this data is also shown in **Extended Data Fig. 1**.

**Extended Data Fig. 3:**
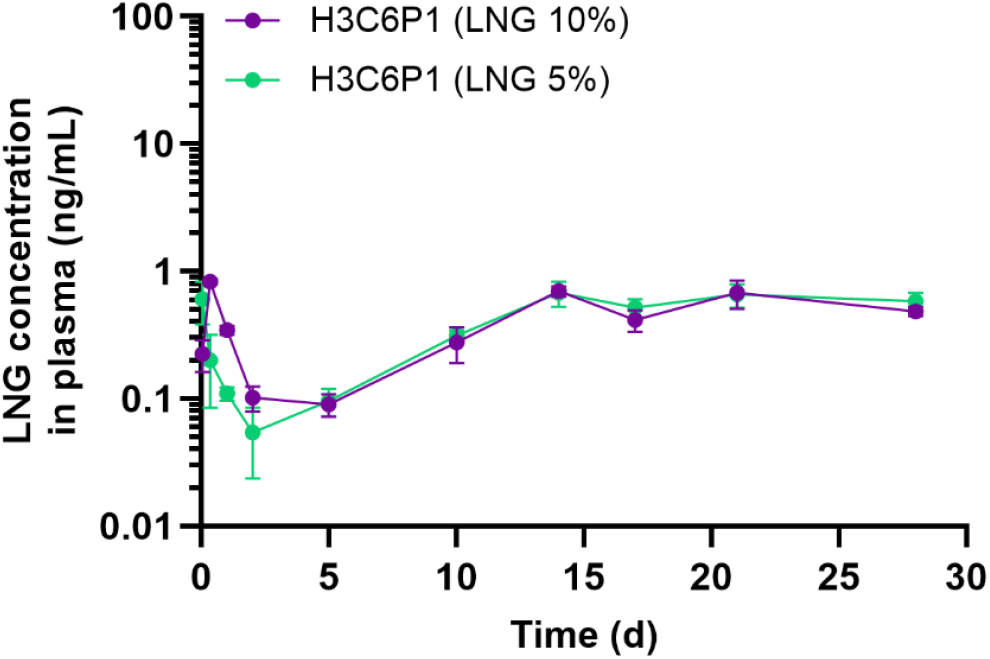
LNG concentration in rat plasma from 5% (w/w) (n = 5) and 10% (w/w) (n = 4) LNG-loaded POE-H3C6P1 MP/HG composite shown as a function of time. A subset of this data is also shown in **Supplementary Fig. 8**.

