## Supplementary Information for "An injectable soft implant for long-acting, reversible, ultra-stable release of therapeutics"

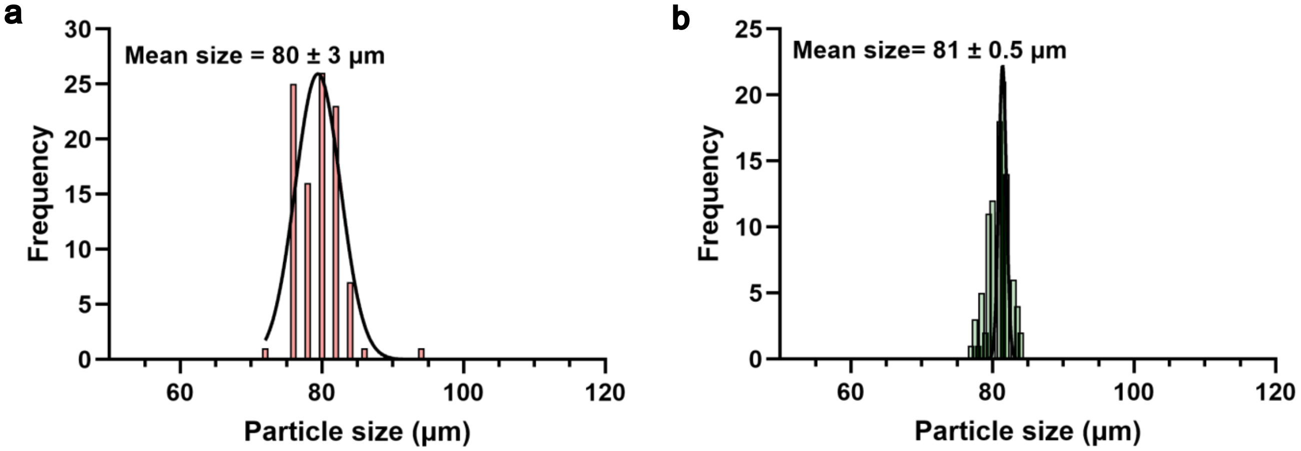


**Supplementary Fig. 1: Histogram of particle size distribution of polymeric microparticles obtained by scanning electron microscopy image analysis (n = 100 particles).** Frequency represents the number of particles counted in each size bin, plotted against particle diameter (µm). **a,** Mean particle size of POE-H3C6P1 MPs was 80 ± 3 µm. **b,** Mean particle size of PLGA (Resomer® RG 503 H) MPs was 81 ± 0.5 µm. See Fig. 2b and f respectively in the main text for the related SEM images.

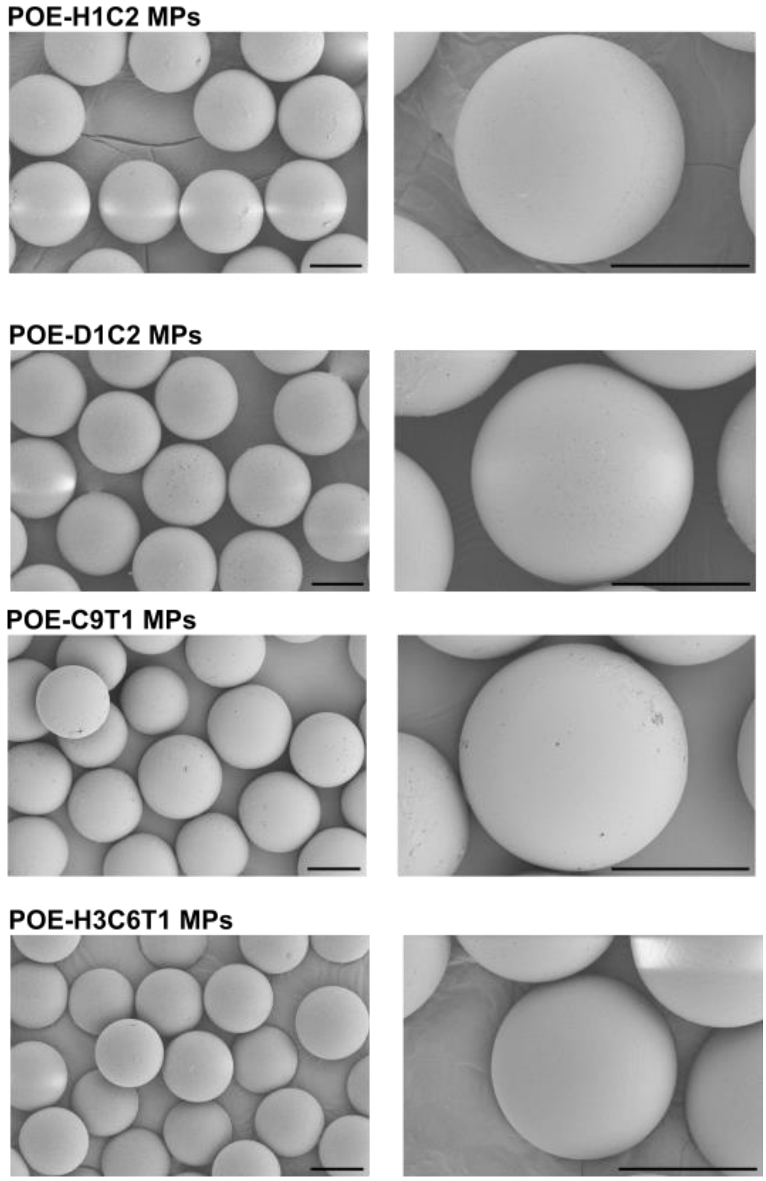


**Supplementary Fig. 2: SEM images of different POE MPs. Scale bar: 50 μm.**

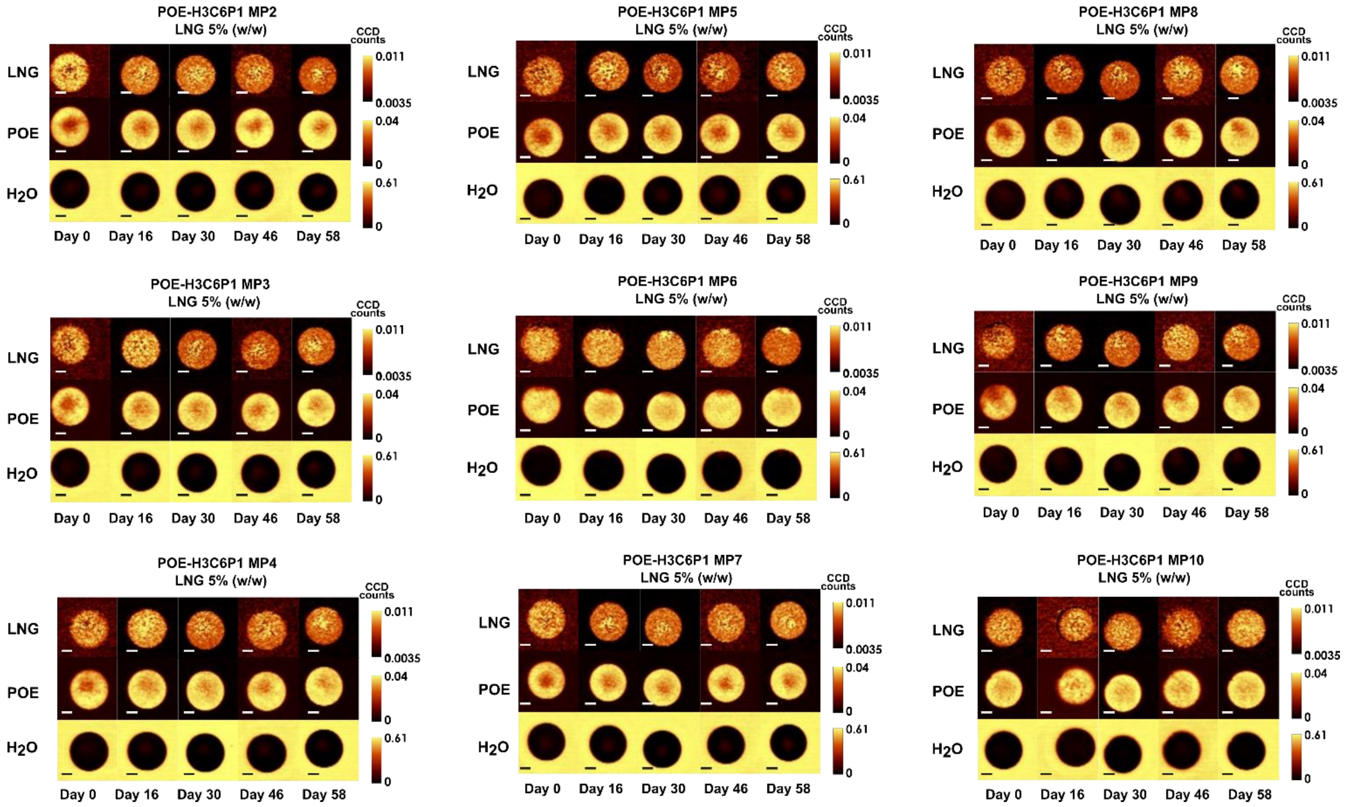


**Supplementary Fig. 3: Raman confocal imaging of single POE-H3C6P1 5% (w/w) LNG loading.** Each individual panel represents a different particle imaged on different days. Scale bar: 20 µm.

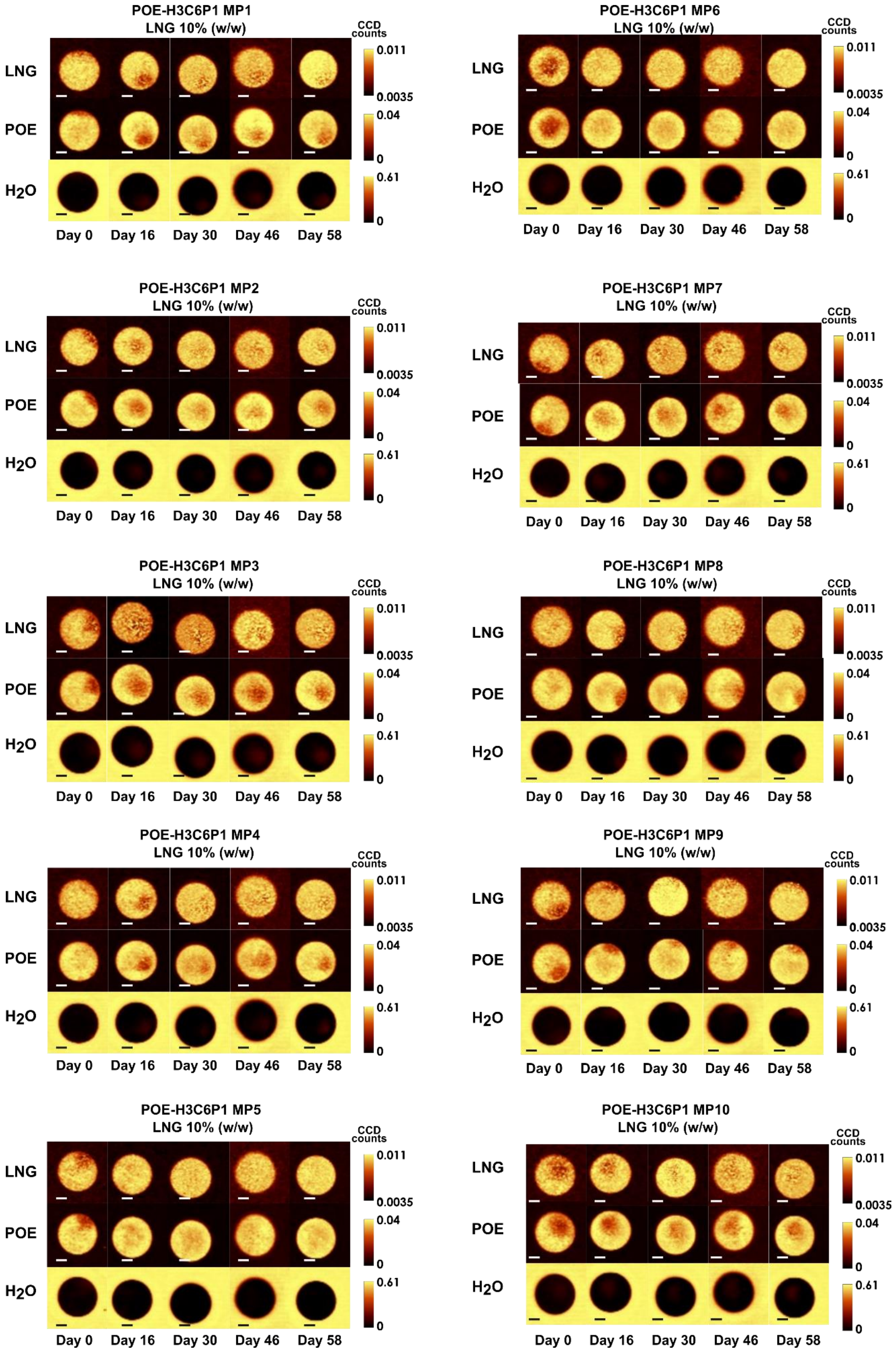


**Supplementary Fig. 4: Raman confocal imaging of single POE-H3C6P1 10% (w/w) LNG loading.** Each individual panel represents a different particle imaged on different days. Scale bar: 20 μm.

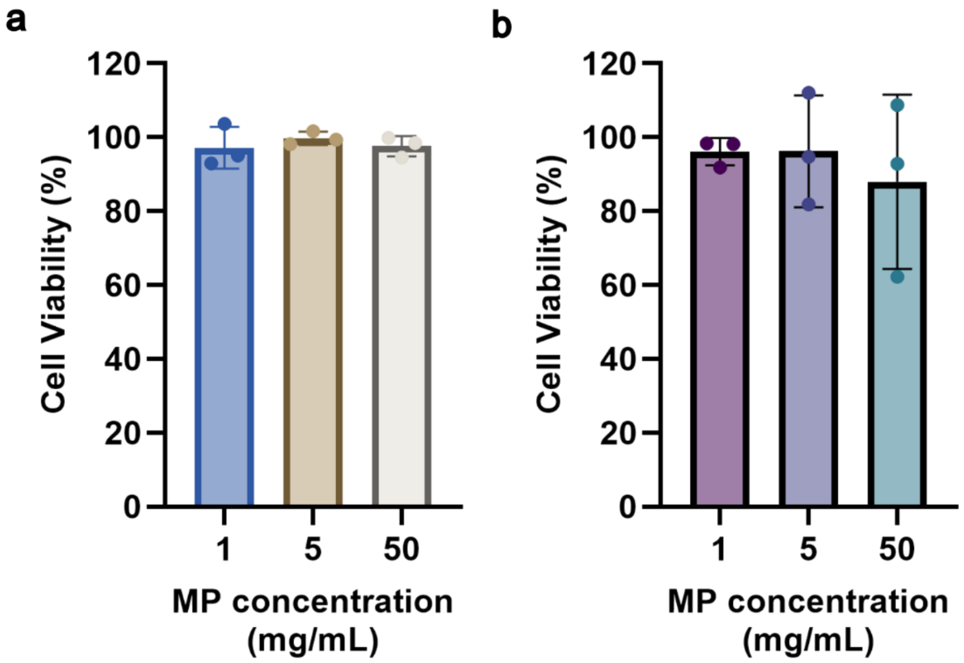


**Supplementary Fig. 5: Cell viability of fibroblast cells following exposure to POE-H3C6P1 microparticles (MPs) at varying concentrations, measured using the AlamarBlue assay. a,** Cells were treated with fresh (non-degraded) POE-H3C6P1 MPs. Data are presented as mean ± SD (N = 3, n = 3 per condition). **b**, POE-H3C6P1 MPs were pre-incubated at 37 °C for 30 days prior to exposure (degraded MPs). Data are presented as mean ± SD (N = 3, n = 3 per condition).

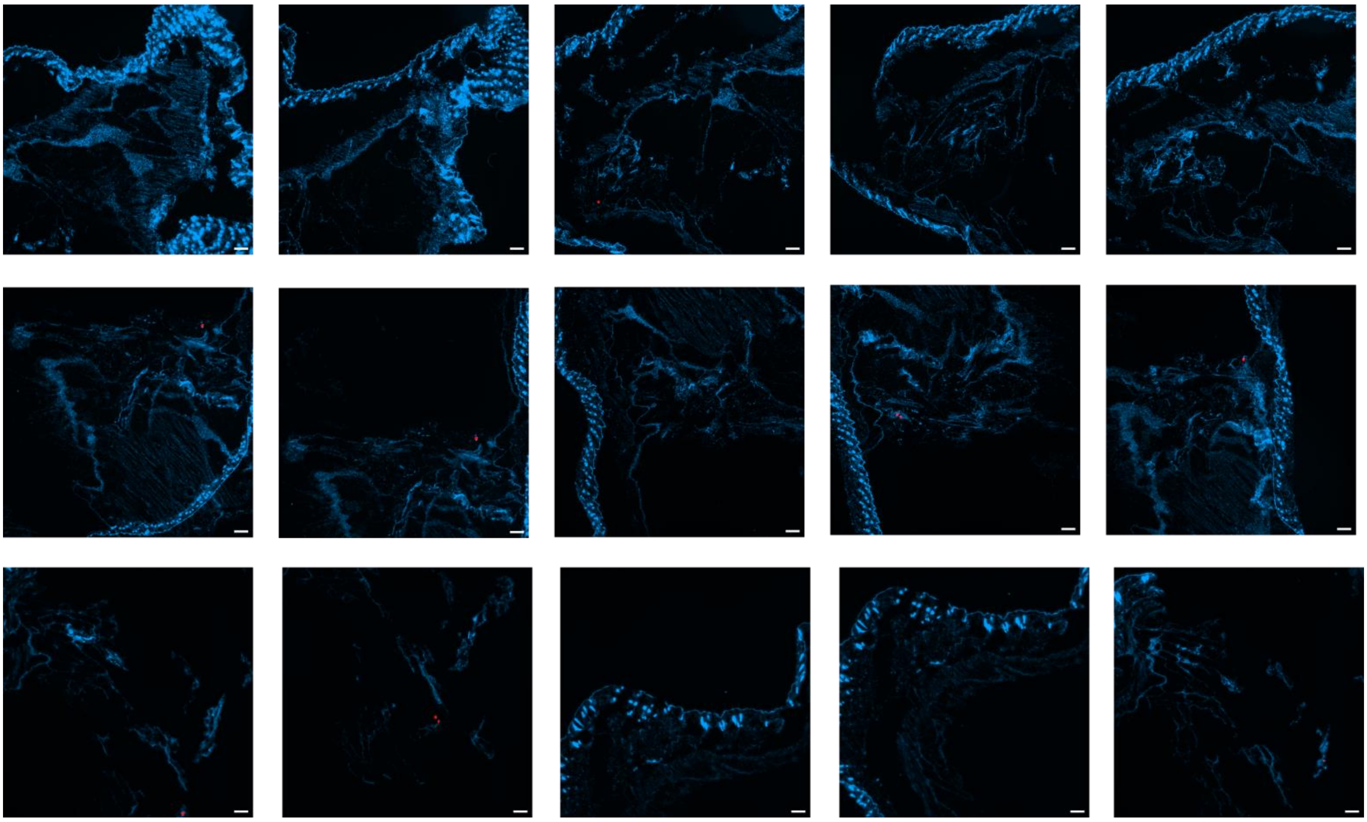

**Supplementary Fig. 6: Fluorescence microscopy images of tissue sections post-removal. Scale bar: 300 µm.** Serial sections of the subcutaneous space after the gel removal through a small incision demonstrate how the vast majority of the PLGA particles (Rhodamine-labelled) is successfully removed.

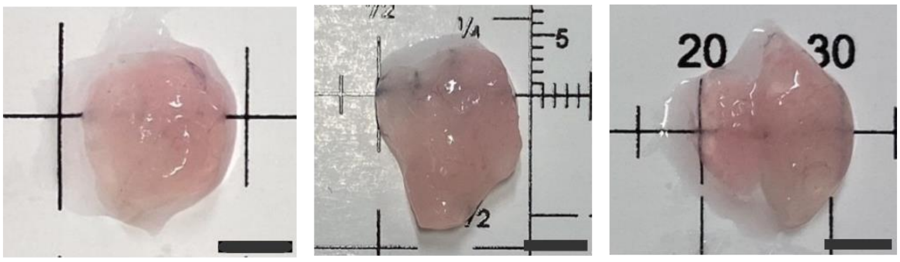


**Supplementary Fig. 7: Images of extracted gels post-removal after 60 days of implantation in Balb/c mice.** Gels removed via small incision display a clear pink colour due to the presence of Rhodamine-labelled PLGA microparticles, confirming how the majority of the microparticles is retained in the gel and can be removed. Scale bar: 5 mm.

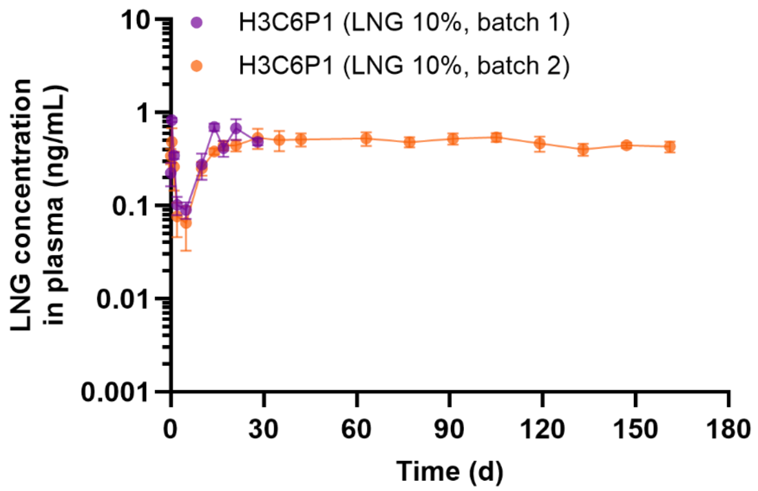


**Supplementary Fig. 8: LNG concentration in rat plasma from 10% (w/w)) LNG-loaded POE-H3C6P1 MP/HG composite shown as a function of time (n = 4 for 28-day study and n = 5 for 161-day study).** A subset of this data is also shown in **Extended Data Fig. 3**.

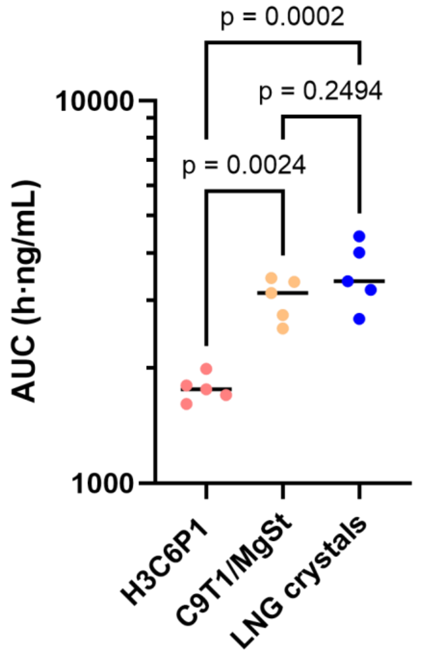


**Supplementary Fig. 9: AUC at the end of the 161-day PK study for the formulations tested.** The measured AUC indicates how same total dose was released from LNG crystals and C9T1/MgSt formulations after 24 weeks, while only ca. 50% of the total dose is released from the H3C6P1 formulation in 24 weeks. One-way ANOVA with Tukey's multiple comparisons post hoc test was used for statistical analysis (n = 5).

**Supplementary Table 1: Description of the groups tested in the pilot 28-day PK study.**

| Formulation | LNG dose (mg) | Particles mass (mg) | Implant volume (mL) | Number of animals |
|---|---|---|---|---|
| POE H3C6P1 (LNG 10%)/HG | 9.1 | 100 | 0.5 | 4 |
| POE H3C6P1 (LNG 5%)/HG | 9.5 | 200 | 1 | 5 |

**Supplementary Table 2: Description of the groups tested in the 161-day PK study. Each group consisted of 5 animals.**

| Formulation | LNG dose (mg) | Particles mass (mg) | Implant volume (mL) |
|---|---|---|---|
| POE H3C6P1/HG | 9.1 | 100 | 0.5 |
| POE C9T1/MgSt/HG | 9.3 | 200 | 1 |
| LNG crystals/HG | 9 | N/A | 0.5 |

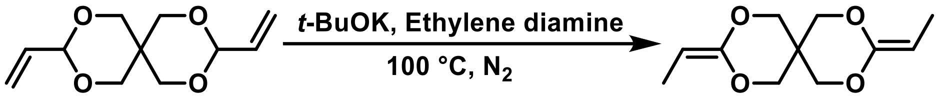


**Supplementary Scheme 1**: Chemical synthesis of the DETOSU monomer from DVTOSU under alkaline conditions.

$^1H$ NMR (400 MHz, $CDCl_3$) δ 4.06 (q, *J* = 6.9 Hz, 2H, =C(***H***)), 3.92 (d, *J* = 1.4 Hz, 4H, O-C($\boldsymbol{H}_2$)-C), 3.85 (d, *J* = 1.4 Hz, 4H, O-C($\boldsymbol{H}_2$)-C), 1.49 (d, *J* = 6.9 Hz, 6H, -C($\boldsymbol{H}_3$)).

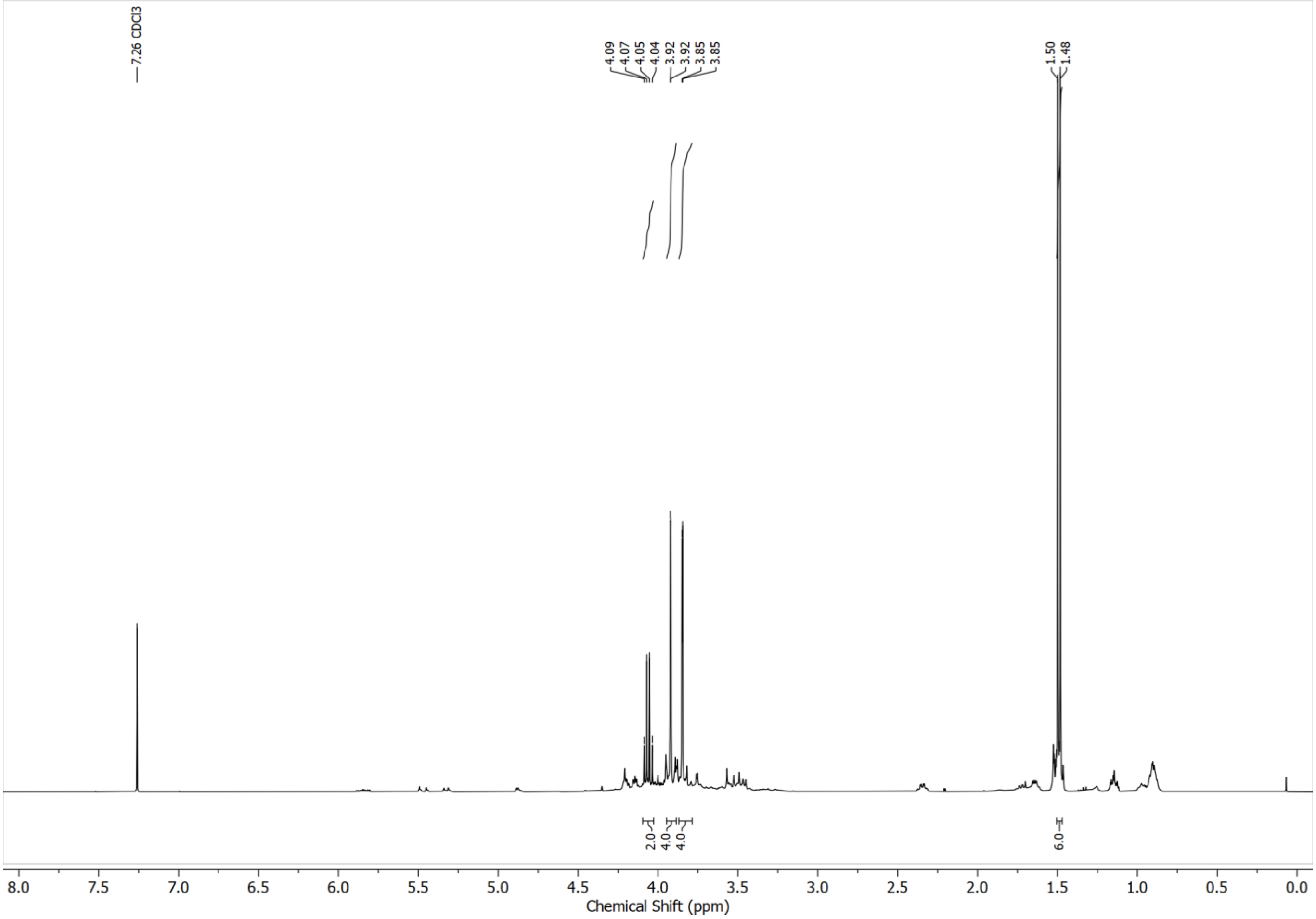


**Supplementary Fig. 10: $^1$H-NMR spectrum of the DETOSU monomer recorded in $CDCl_3$ at 400 MHz at 25 °C.**

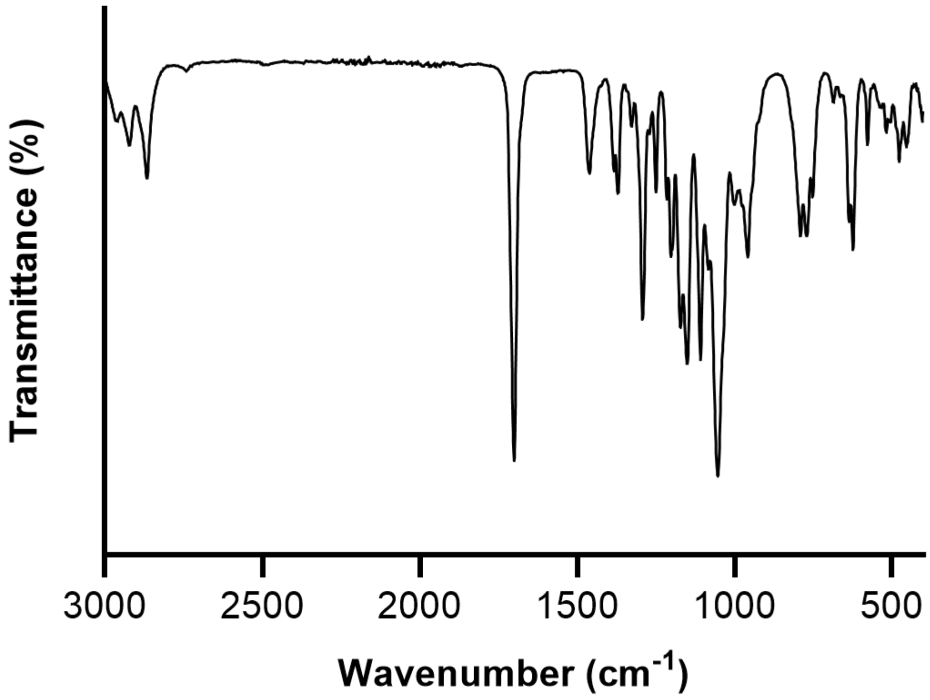


**Supplementary Fig. 11: FTIR spectrum of DETOSU monomer recorded using ATR-FTIR.**

**Polymer NMRs**

**Note:** The $^{1}$H NMR spectra exhibit broad multiplets, minor resonances in the 6.0–5.0 ppm region, and non-integral signal intensities. These features are expected and attributable to the random monomer sequence distribution in block copolymers, the dispersity of polymer chain lengths, variations in chain-termination end groups (e.g., –OH from alcohol termination or $=CH_2$ from DETOSU back-isomerisation), and the fractional stoichiometric incorporation of diol units during POE synthesis. The following assignments correspond to the expected $^{1}$H spectra arrising from an average POE 'block'.

$^{1}$H NMR (400 MHz, $CDCl_3$) δ 6.17-6.12 (m, 0.1H, =C(***H***$_2$)), 5.64-5.60 (m, 0.1H, =C(***H***$_2$)), 4.15-3.20 (m, 12.1H, O-C(***H***$_2$)-C / O-C(***H***)-C), 1.87-1.03 (m, 13H, -C(***H***$_2$)-), 0.91 (m, 6.3H, -C(***H***$_3$)).

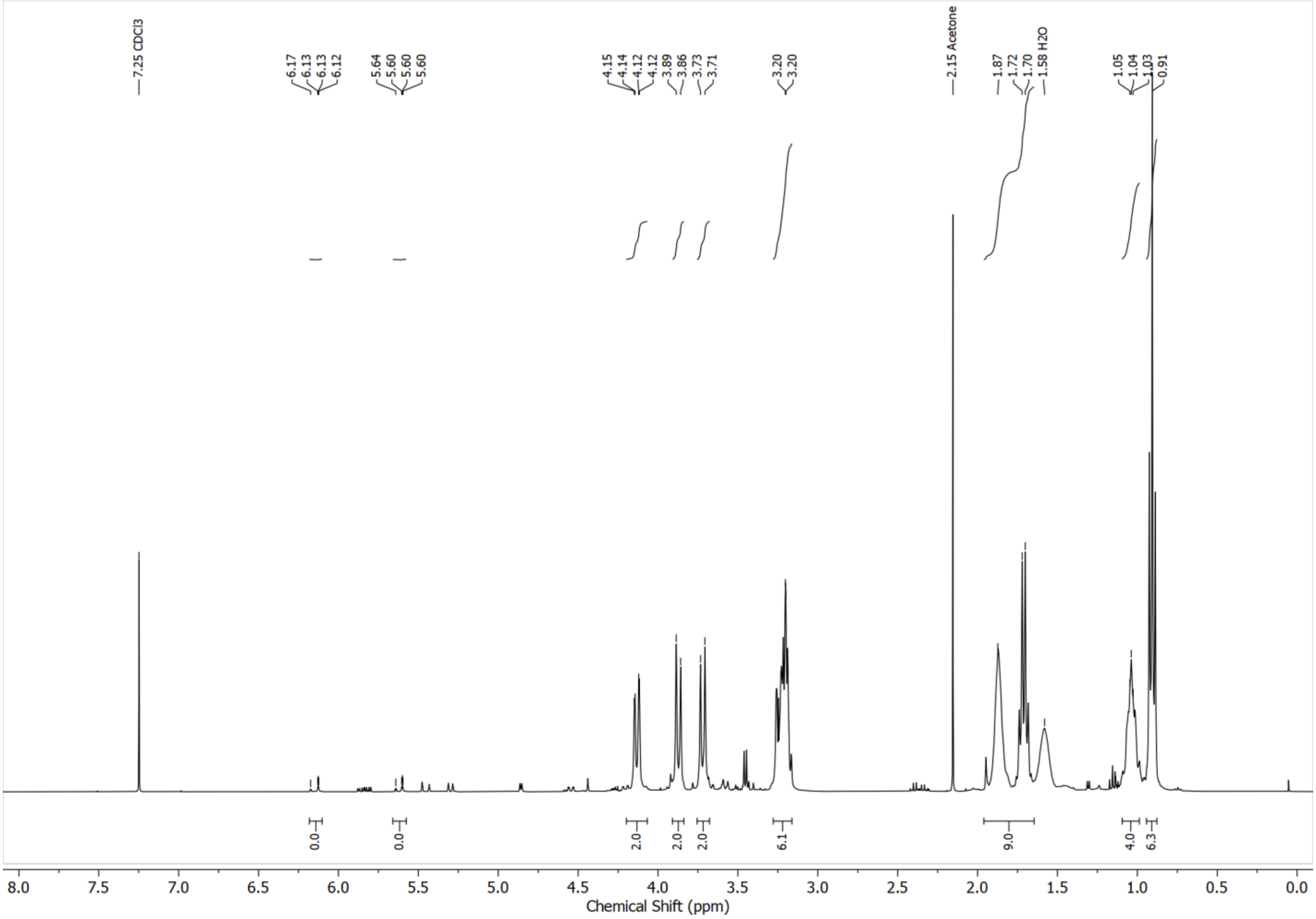


**Supplementary Fig. 12: $^{1}$H-NMR spectrum of POE-C9G1 recorded in $CDCl_3$ at 400 MHz.**

$^1H$ NMR (400 MHz, $CDCl_3$) δ 4.18-3.28 (m, 12H, O-C(**$H_2$**)-C), 1.87-1.46 (m, 12H, -C(***$H_2$***)-), 0.95 (m, 6H, -C(***$H_3$***)).

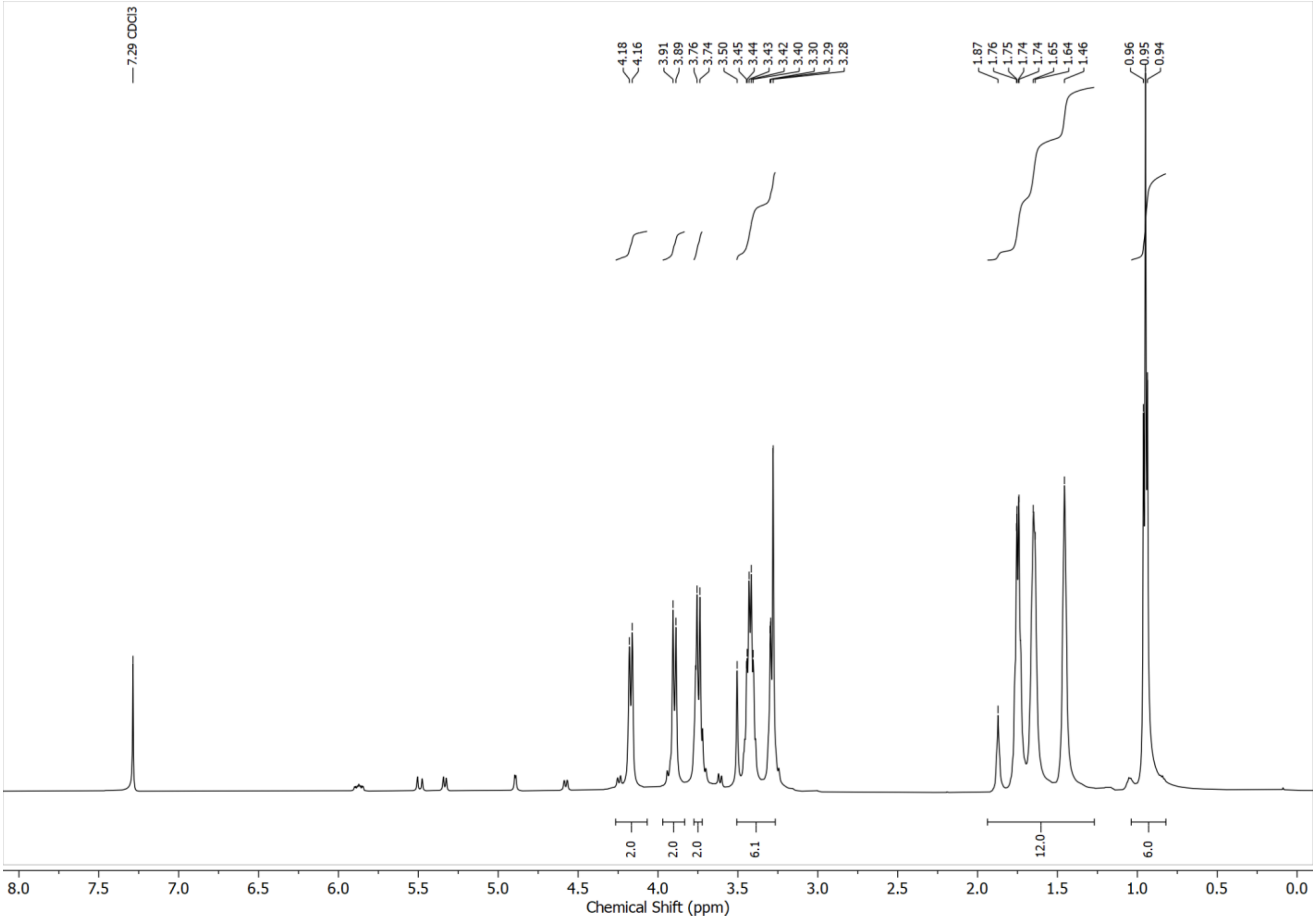


**Supplementary Fig. 13: $^1H$-NMR spectrum of POE-H recorded in $CDCl_3$ at 400 MHz.**

$^{1}H$ NMR (400 MHz, $CDCl_3$) δ 4.14-3.18 (m, 12H, O-C($\mathbf{H_2}$)-C), 1.87-1.46 (m, 14H, -C($\boldsymbol{H}_2$)- / -C($\boldsymbol{H}$)-), 0.90 (m, 6H, -C($\boldsymbol{H}_3$)).

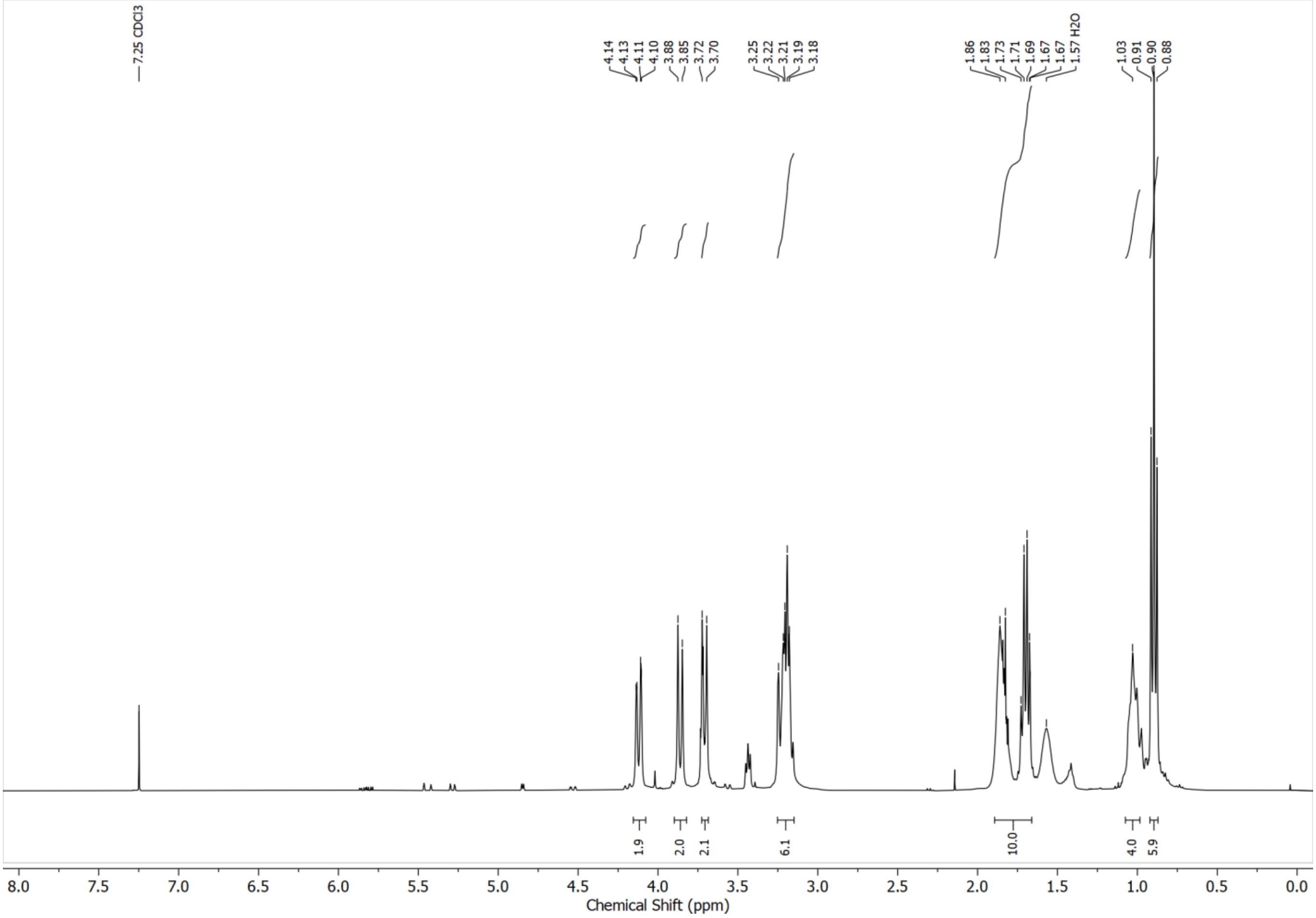


**Supplementary Fig. 14: $^{1}$H-NMR spectrum of POE-C recorded in $CDCl_3$ at 400 MHz.**

$^1H$ NMR (400 MHz, $CDCl_3$) δ 4.14-3.19 (m, 12H, O-C(**$H_2$**)-C), 1.71-1.03 (m, 13.3H, -C(***$H_2$***)- / -C(***H***)-), 0.91 (m, 6H, -C(***$H_3$***)).

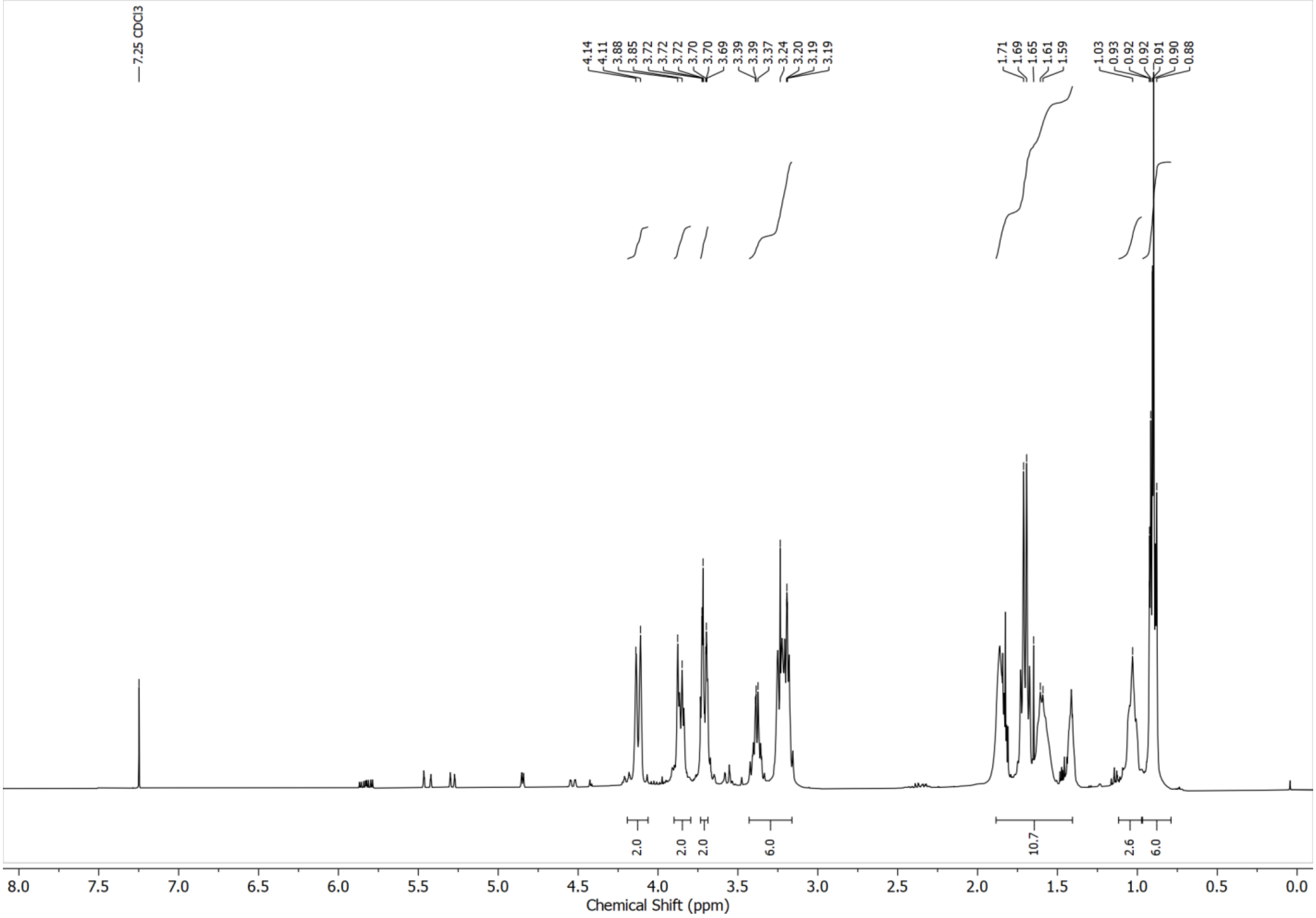


**Supplementary Fig. 15: $^1$H-NMR spectrum of POE-H1C2 recorded in $CDCl_3$ at 400 MHz.**

$^{1}H$ NMR (400 MHz, $CDCl_3$) δ 4.11-3.19 (m, 12H, O-C(**H₂**)-C), 1.86-1.03 (m, 16H, -C(***H***₂)- / -C(***H***)-), 0.90 (m, 6H, -C(***H***₃)).

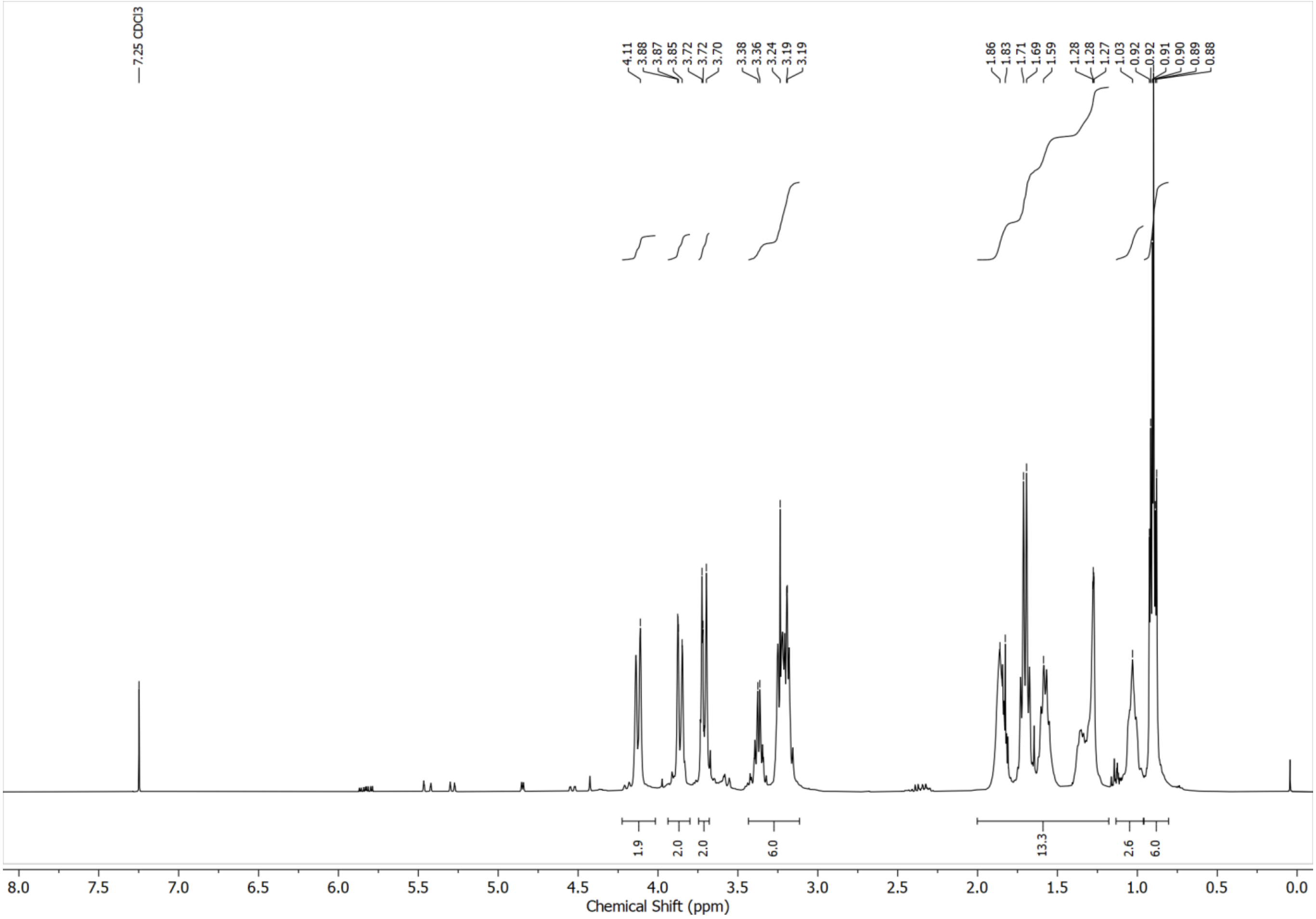

**Supplementary Fig. 16: $^{1}H$-NMR spectrum of POE-D1C2 recorded in $CDCl_3$ at 400 MHz.**

$^{1}H$ NMR (400 MHz, $CDCl_3$) δ 4.14-3.19 (m, 12.8H, O-C(**$H_2$**)-C), 1.88-1.03 (m, 13H, -C(***$H_2$***)- / -C(***H***)-), 0.90 (m, 6H, -C(***$H_3$***)).

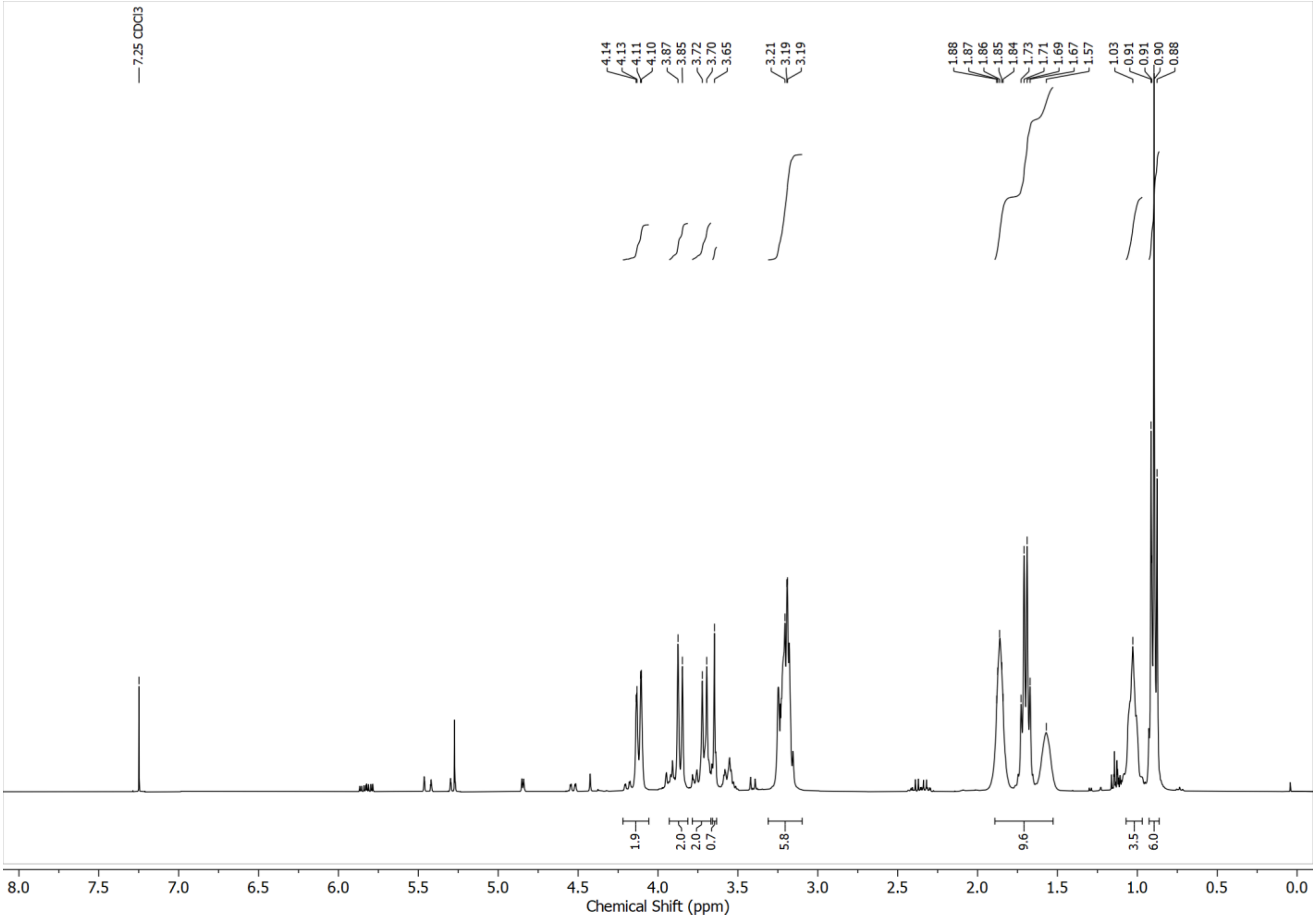


**Supplementary Fig. 17: $^{1}H$-NMR spectrum of POE-C9T1 recorded in $CDCl_3$ at 400 MHz.**

$^{1}$H NMR (400 MHz, $CDCl_3$) δ 4.04-3.15 (m, 12H, O-C($\mathbf{H_2}$)-C), 2.58-2.56 (m, 0.8H, N-C($\mathbf{H_2}$)-C), 1.85-1.01 (m, 12.4H, -C($\boldsymbol{H}_2$)- / -C($\boldsymbol{H}$)-), 0.90 (m, 6H, -C($\boldsymbol{H}_3$)).

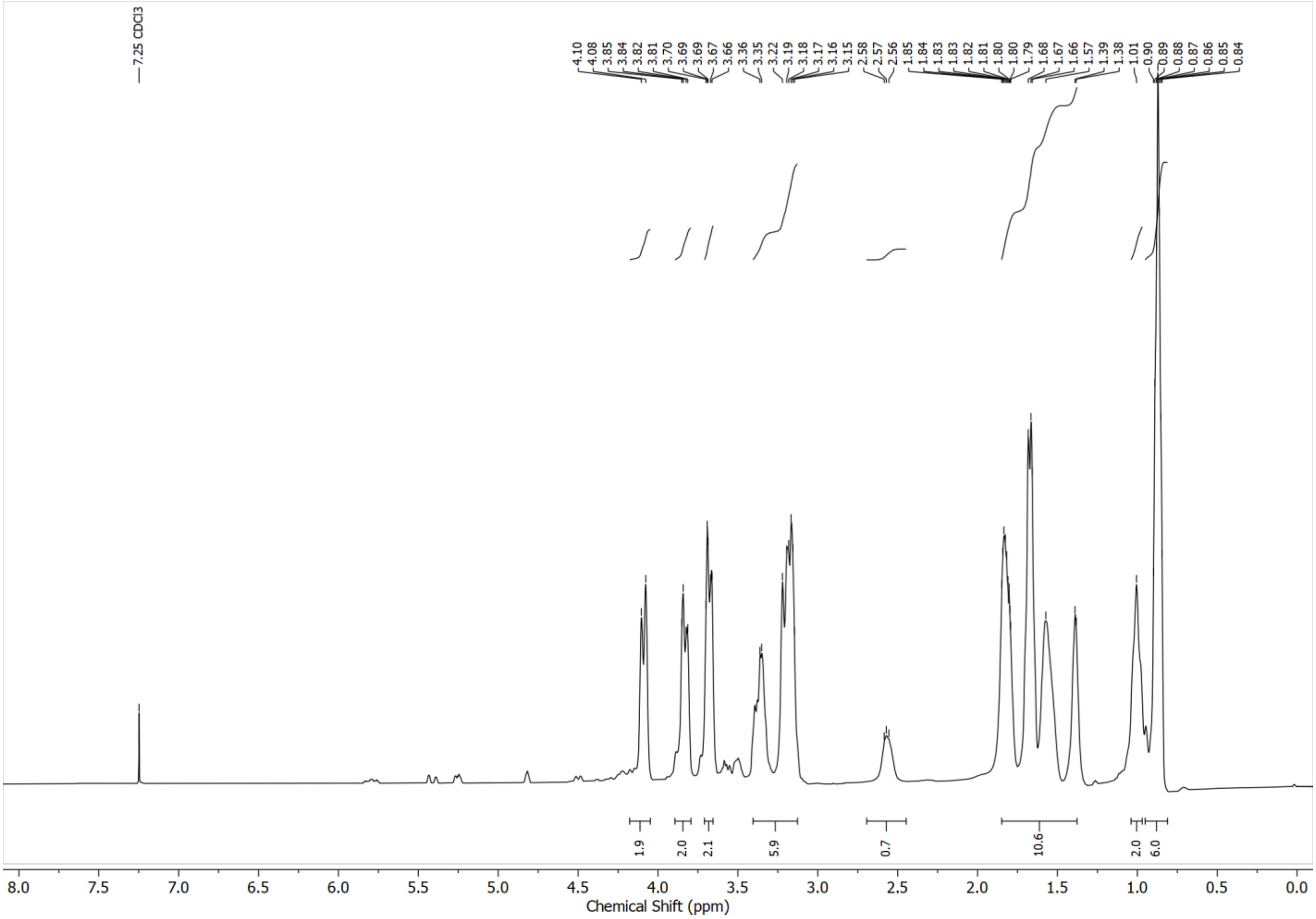


**Supplementary Fig. 18: $^{1}$H-NMR spectrum of POE-H3C6P1 recorded in $CDCl_3$ at 400 MHz.**

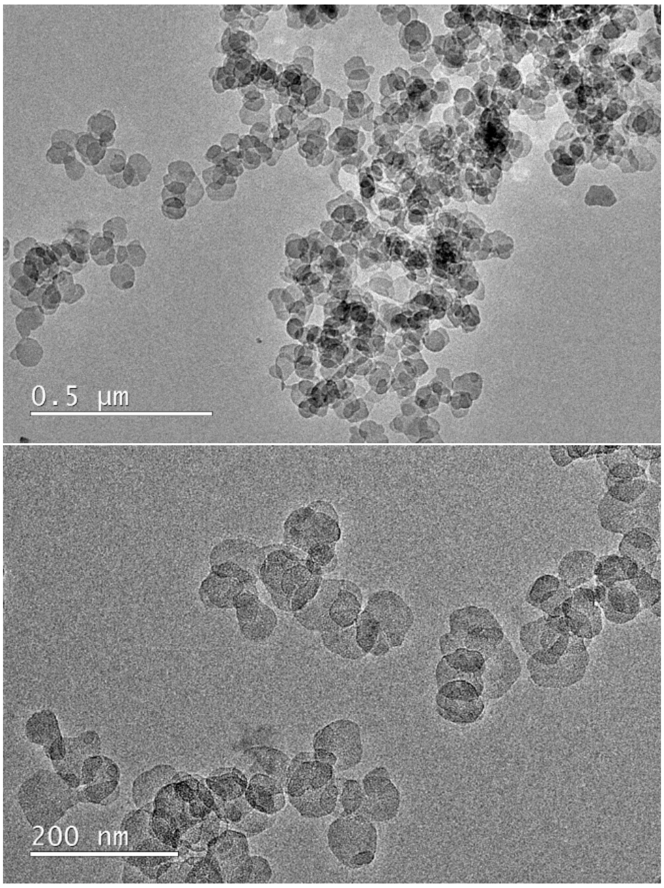


**Supplementary Fig. 19: TEM images of $Mg(OH)_2$ nanoparticles.**
